# Multiplexed spatial mapping of chromatin features, transcriptome, and proteins in tissues

**DOI:** 10.1101/2024.09.13.612892

**Authors:** Pengfei Guo, Liran Mao, Yufan Chen, Chin Nien Lee, Angelysia Cardilla, Mingyao Li, Marek Bartosovic, Yanxiang Deng

## Abstract

The phenotypic and functional states of a cell are modulated by a complex interactive molecular hierarchy of multiple omics layers, involving the genome, epigenome, transcriptome, proteome, and metabolome. Spatial omics approaches have enabled the capture of information from different molecular layers directly in the tissue context. However, current technologies are limited to map one to two modalities at the same time, providing an incomplete representation of cellular identity. Such data is inadequate to fully understand complex biological systems and their underlying regulatory mechanisms. Here we present spatial-Mux-seq, a multi-modal spatial technology that allows simultaneous profiling of five different modalities, including genome-wide profiles of two histone modifications and open chromatin, whole transcriptome, and a panel of proteins at tissue scale and cellular level in a spatially resolved manner. We applied this technology to generate multi-modal tissue maps in mouse embryos and mouse brains, which discriminated more cell types and states than unimodal data. We investigated the spatiotemporal relationship between histone modifications, chromatin accessibility, gene and protein expression in neuron differentiation revealing the relationship between tissue organization, function, and gene regulatory networks. We were able to identify a radial glia spatial niche and revealed spatially changing gradient of epigenetic signals in this region. Moreover, we revealed previously unappreciated involvement of repressive histone marks in the mouse hippocampus. Collectively, the spatial multi-omics approach heralds a new era for characterizing tissue and cellular heterogeneity that single modality studies alone could not reveal.

## Main

The intricate interplay between genotype and phenotype is shaped by a molecular hierarchy spanning multiple omics layers, involving the genome, epigenome, transcriptome, proteome, and metabolome^1–3^. In addition, the organization of cellular compartments, structures, and intercellular interactions is critical to the functional state of a cell in multicellular organisms^3^. Therefore, methodological and technological advances that allow simultaneous measurement of different layers of molecular information from cells within their native tissue context are crucial^1^. Recent advancements in multi-modal spatial omics have aided in resolving biological complexity by studying different molecular analytes within their original tissue contexts^4–8^. For example, parallel epigenomic profiling with gene expression uncovered new information of epigenetic priming, differentiation, and gene regulation within the tissue architecture^4,5^. Spatial co-mapping of the whole transcriptome and a panel of proteins substantially improved cell clustering and enhanced the discovery power across tissue regions, compared with unimodal measurements^6–8^. However, experimental integration of all these modalities is lacking, providing an incomplete representation of cellular states, and is inadequate to develop a fundamental understanding of the complex biological systems and their underlying regulatory mechanisms. In addition, cellular transcription programs are determined through the action of multiple epigenetic modalities, including transcription factors, and co-occurrence of synergistic or antagonistic histone marks^9^. The effects of these interactive chromatin regulatory factors on downstream gene or protein expression are missing from current single cell and spatial approaches.

In this study, we report a multi-modal spatial technology that allows simultaneous profiling of up to five different modalities, including open chromatin and two histone modifications, whole transcriptome, and a panel of proteins at tissue scale and cellular level in a spatially resolved manner. This was achieved by integrating microfluidic *in situ* barcoding^4,7,10,11^ and the nanobody-tethered transposition chemistry directly in tissue followed by high-throughput Next-Generation Sequencing (NGS)^9,12^. We applied this new technology to generate multi-modal tissue maps in mouse embryos and mouse brains, which enabled investigation of the intermolecular dynamics among chromatin states characterized by combinations of epigenetic factors, gene and/or protein expression, and tissue development, in a spatially resolved manner.

### Technology workflow

The workflow of simultaneous profiling of chromatin accessibility and/or a panel of cell surface proteins with two histone modifications and gene expression on the same tissue cryosections is shown schematically in Fig. 1a and Extended Data Fig. 1. A frozen tissue section was first fixed with formaldehyde and *in situ* Tn5 transposition was performed to insert a unique barcoded ligation linker to the accessible DNA loci. The same tissue section was then incubated with two primary antibodies targeting different histone modifications simultaneously, such as the combination of H3K27me3 with H3K27ac or H3K4me3. Afterwards, species specific nanobody-Tn5 fusion proteins loaded with unique barcoded ligation linkers were added to enable the demultiplexing of different histone modification loci. For co-profiling of proteins, the fixed frozen tissue section was stained with a panel of poly-A-tailed oligo-conjugated antibodies, which recognize surface antigens. Next, *in situ* reverse transcription was performed using the biotinylated poly-T RT primer to capture both oligo-conjugated antibodies and mRNA. Next, barcodes A (A_1_-A_50_ or A_1_-A_100_) and barcodes B (B_1_-B_50_ or B_1_-B_100_) were sequentially flowed over the tissue using microchannels and were ligated to the universal ligation liker, which formed a two-dimensional grid of spatially barcoded tissue pixels (n =2,500 or 10,000), allowing all of modalities from the same pixel share the same spatial barcodes. Finally, barcoded complementary DNA (cDNA) and genomic DNA (gDNA) fragments were released by reverse crosslinking. cDNAs were separated from gDNA by streptavidin beads. Sequencing libraries for cDNAs and gDNA were then separately constructed. The protein library and mRNA library can be further separated by SPRI bead.

**Fig. 1.**
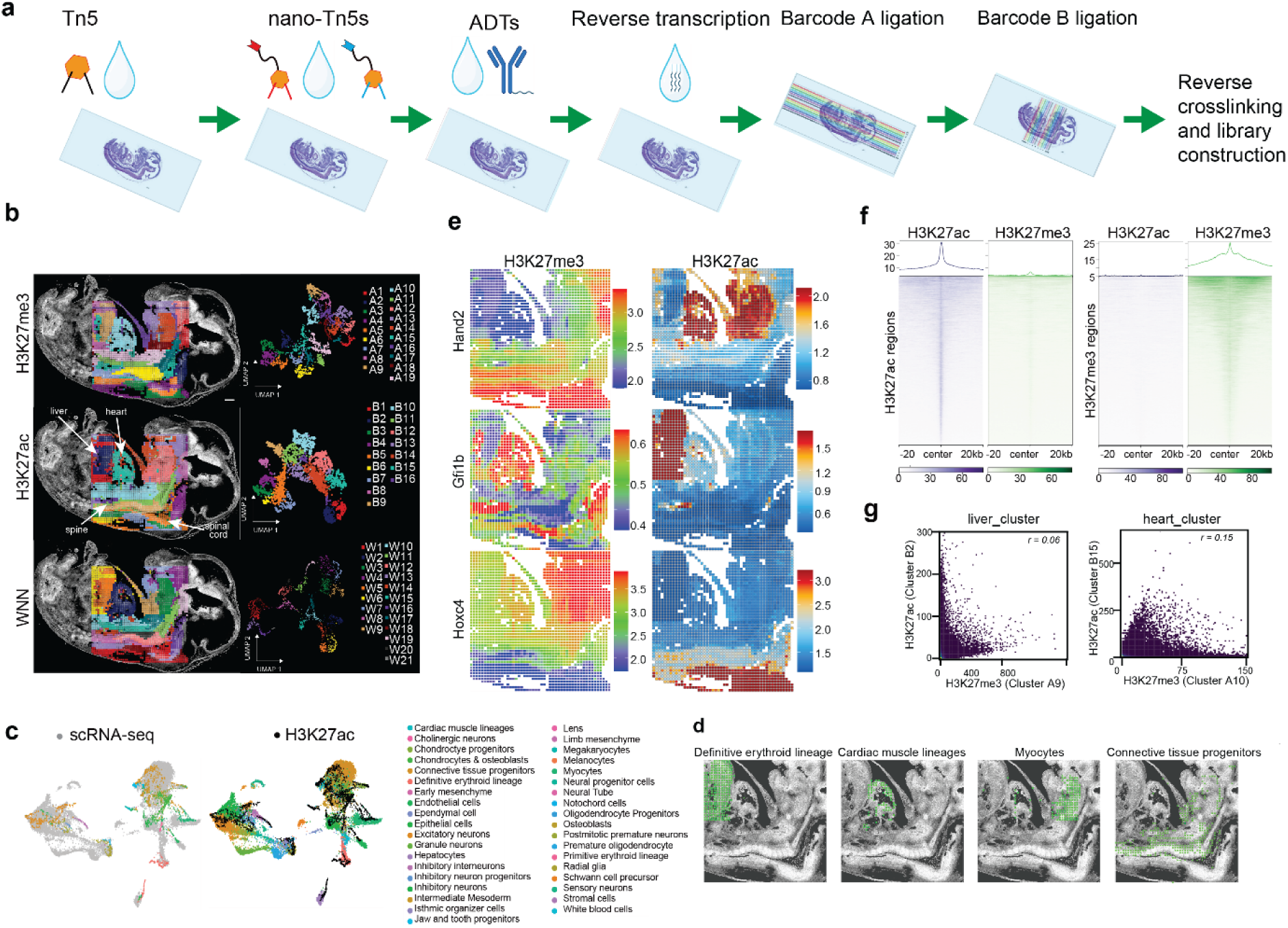
Spatial-Mux-seq co-profiling of H3K27me3 and H3K27ac modifications in E13 mouse embryos with integrative analysis. Sample: E13_50_μm_1. **a,** A schematic overview illustrating the workflow for spatial multimodal profiling of chromatin modifications at the tissue scale. The workflow starts with tissue fixation, followed by Tn5 and nano-Tn5 transposition, antibody-derived tags (ADTs) application, reverse transcription, and sequential ligation of barcodes A and B. The process is finalized with reverse crosslinking and library construction to enable comprehensive spatial analysis. **b,** Spatial distribution and Uniform Manifold Approximation and Projection (UMAP) embeddings derived from unsupervised clustering analysis of H3K27me3 and H3K27ac histone modifications. This panel includes an integrated analysis using Weighted Nearest Neighbor (WNN) methodology, displaying the spatially resolved chromatin state across key anatomical regions such as the liver, heart, and spinal cord. **c,** Integration of single-cell RNA sequencing (scRNA-seq) data^14^ with spatial-Mux-seq H3K27ac profiling. The alignment of cell types identified in scRNA-seq (left) with spatially resolved H3K27ac data (middle). The cell types identified through scRNA-seq are listed (right). **d,** Spatial mapping of selected cell types identified through label transfer from scRNA-seq to spatial-Mux-seq data. **e,** Spatial mapping of key developmental marker genes, showing their corresponding H3K27me3 and H3K27ac histone modifications. **f,** Metagene plots showing the distribution of H3K27me3 and H3K27ac in fetal liver clusters obtained by spatial-Mux-seq around specific H3K27me3 and H3K27ac peaks. The peaks were defined from ENCODE datasets. **g,** Scatter plots showing correlation of H3K27me3 and H3K27ac signal in the liver and heart clusters. The peaks were defined from ENCODE datasets. *r*, Pearson correlation coefficient. scale bar: 500 μm.

### Evaluation of spatial-Mux-seq profiling of two histone modifications

Nanobody-based multimodal CUT&Tag has not previously been used directly in tissues. Therefore, we initially assessed the specificity of *in situ* transposition using species-specific nanobody-Tn5 fusion proteins. We targeted two distinct histone modifications on the same tissue section: H3K27me3 (trimethylation of lysine 27 on histone H3), a repressive mark typically found at silenced genes, is mediated by the Polycomb Repressive Complex 2 (PRC2) and plays a crucial role in maintaining gene repression during development and preserving cell identity. H3K27ac (acetylation of lysine 27 on histone H3) marks active enhancers and promoters is associated with active gene expression. These two histone modifications, H3K27me3 and H3K27ac, are mutually exclusive and represent opposing chromatin states, making them an ideal model for evaluating the specificity of the nanobody-based *in situ* transposition method.

We first benchmarked spatial-Mux-seq in E13 sagittal mouse embryo sections with 50-µm resolution (E13_50_µm_1), and obtained a median of 17,677 and 9,893 unique fragments per pixel for H3K27me3 and H3K27ac respectively (Extended Data Fig. 2a-b). We benchmarked these matrics by comparing them to individual omics datasets previously obtained using spatial-CUT&Tag^11^. We found that transcriptional start site (TSS) enrichment scores for both modalities closely aligned with those obtained in other single-modality studies (Extended Data Fig. 2c). Notably, our method obtained comparable number of unique fragments per pixel, matching the performance of single-modality profiling (Extended Data Fig. 2c)^11^. Reproducibility across replicates from different experiments (E13_50_µm_1 and E13_50_µm_2) was high (Extended Data Fig. 3a-c), as demonstrated by Pearson correlation of *r* = 0.93 for H3K27me3 and *r* = 0.91 for H3K27ac (Extended Data Fig. 3d). Additionally, consistent and reproducible peaks were obtained across replicates (Extended Data Fig. 3e), and the insert size distributions of the co-profiled histone modifications (H3K27me3/H3K27ac) showed expected and typical nucleosomal phasing pattern (Extended Data Fig. 3f). These results demonstrated the robustness of our method.

We then performed unsupervised clustering and identified 19 and 16 clusters for H3K27me3 (An) and H3K27ac (Bn) respectively (Fig. 1b). Both exhibited distinct spatial patterns consistent with the tissue histology of an adjacent section stained with hematoxylin and eosin (Extended Data Fig. 2d). For example, Cluster A10 of H3K27me3 and cluster B15 of H3K27ac corresponded to the embryonic heart; Cluster A9 of H3K27me3 and cluster B2 of H3K27ac were the liver; Cluster A6 of H3K27me3 and cluster B9 of H3K27ac located in the spine region. To integrate both modalities, weighted nearest neighbor (WNN) analysis^13^ was used, resulting in improved clustering in the low-dimensional embedding (Fig. 1b). The alluvial diagram and UMAP projection further illustrated that the WNN clustering effectively recapitulated and refined clusters identified by H3K27ac and H3K27me3 (Extended Data Fig. 4a-b). Cell types for each cluster were then assigned by label transfer from mouse embryonic (E13.5) scRNA-seq data^14^ to spatial-Mux-seq data (H3K27ac) (Fig. 1c). For instance, definitive erythroid cells appeared predominantly in the liver, cardiac muscle lineages were identified within the heart region, myocytes were enriched in both skeletal muscles and the heart region, and connective tissue progenitors distributed across various regions where connective tissues are developing (Fig. 1d).

The development of the mouse embryo is an intricate and highly regulated process that involves the coordinated expression and silencing of numerous genes^15^. We then explored the spatial patterns of specific marker genes and examined the interplay between active (H3K27ac) and repressive (H3K27me3) histone marks (Fig. 1e, Extended Data Fig. 5). For H3K27me3 and H3K27ac, the chromatin silencing score (CSS) and gene activity score (GAS) were calculated to predict the gene expression respectively^16^. *Hand2*, which is an important regulator of craniofacial development and plays an essential role in cardiac morphogenesis^17,18^, showed an enrichment of H3K27ac but lacked H3K27me3 in the jaw and heart region (Fig. 1e). As another example, *Gfi1b*, which is essential for the development of the erythroid and megakaryocytic lineages^19^, showed high GAS of H3K27ac and low CSS of H3K27me3 in the liver region. Similarly, in the liver region, we noted significant enrichment of H3K27ac at *Nprl3* locus (Extended Data Fig. 5a), emphasizing its critical role in the erythroid development^20,21^. In the craniofacial region, there was notable enrichment of H3K27me3, but not H3K27ac, observed at the *Hoxc4* locus (Fig. 1e). Regarding the *Sox2* gene, most clusters exhibited significant enrichment of H3K27me3 except the spinal cord region (Extended Data Fig. 5b), in which *Sox2* is required to maintain the properties of neural progenitor cells within the spinal cord region^22^.

The correlation between epigenetic marks and transcript abundance was further studied by comparing the CSS and GAS with scRNA-seq data^14^. In excitatory neurons, we observed a positive correlation between H3K27ac and gene expression, alongside an anticorrelation with H3K27me3 (Extended Data Fig. 6a-c). Marker genes such as *Ina*, *Crmp1*, and *Atp1a3* exhibited significant enrichment with H3K27ac and minimal enrichment with H3K27me3 in the excitatory neuron region (Extended Data Fig. 6d), highlighting the interplay between active (H3K27ac) and repressive (H3K27me3) histone marks in regulating gene expression.

We then further verified the specificity of each modality by selecting highly specific peaks for H3K27me3 and H3K27ac within the liver region. This analysis revealed significant enrichment of the respective modifications within the corresponding set of marker peaks (Fig. 1f). Additionally, we analyzed H3K27me3/H3K27ac signals within liver and heart clusters, finding no significant correlations between these histone marks (Fig. 1g). These results collectively demonstrated the specificity and efficacy of spatial-Mux-seq in profiling multiple histone modifications in the same tissue section.

### Spatial four-modal profiling of multiple epigenetic modalities and transcriptome

Single cell nanobody-based CUT&Tag has been employed for co-measurement of open chromatin^9^ or cell surface markers^12^, while leave the transcriptome unexplored. To address this limitation and study the intermolecular dynamics between multiple epigenetic regulatory factors and gene and/or protein expression and tissue development, we profiled chromatin accessibility (ATAC), two histone modifications (H3K4me3 and H3K27me3), and transcriptome simultaneously, altogether capturing four molecular layers in the same tissue section at 50-µm resolution (E13_50_µm_3). We obtained a median of 39,014 unique fragments for ATAC, 6,657 for H3K4me3, and 8,496 for H3K27me3 per pixel (Extended Data Fig. 7a-b). These results were benchmarked by comparing with the individual omics data from spatial-CUT&Tag^11^ as well as co-profiled modalities from spatial-ATAC-RNA-seq^4^. Each modality exhibited similar counts of unique fragments (Extended Data Fig. 2i), demonstrating that the inclusion of more modalities does not compromise data quality. Additionally, we observed matched TSS enrichment scores for each modality (Extended Data Fig. 2i). For the RNA portion, a total 22,171 genes were detected with an average of 1,569 genes and 2,538 UMIs per pixel (Extended Data Fig. 7b-c). These results are consistent with RNA results from spatial-ATAC-RNA-seq^4^ performed on the same tissue type. Unsupervised clustering identified, 10 clusters for ATAC (cluster An), 7 clusters for H3K4me3 (cluster Bn), 9 clusters for H3K27me3 (cluster Cn), and 11 clusters for RNA (cluster Rn) (Fig. 2a), which showed concordance in cluster assignment and agreed with tissue histology. For example, the heart region can be identified from different modalities: cluster A4 of ATAC data, cluster B5 of H3K4me3 data, cluster C3 of H3K27me3 data, and cluster R6 of RNA data. While most clusters were identified across all four modalities, we found that few clusters were only revealed by specific molecular layers. For instance, the liver region could be further distinguished into two distinct clusters (A1 and A2) from the ATAC data but not resolved in the H3K4me3 and H3K27me3 data (Fig. 2a), where canonical E2F activator *E2f2* had stronger open chromatin signals in the A2 liver cluster compared with A1 liver cluster (Fig. 2b-c). Additionally, we intersected ATAC, H3K4me3, and H3K27me3 peaks from the liver cluster, and observed that H3K4me3 and ATAC peaks showed strong overlap (8,324 overlapping regions), and a subset of genomic regions demonstrated variability in all three modalities simultaneously (4,165 overlapping regions) (Extended Data Fig. 7d).

**Fig. 2.**
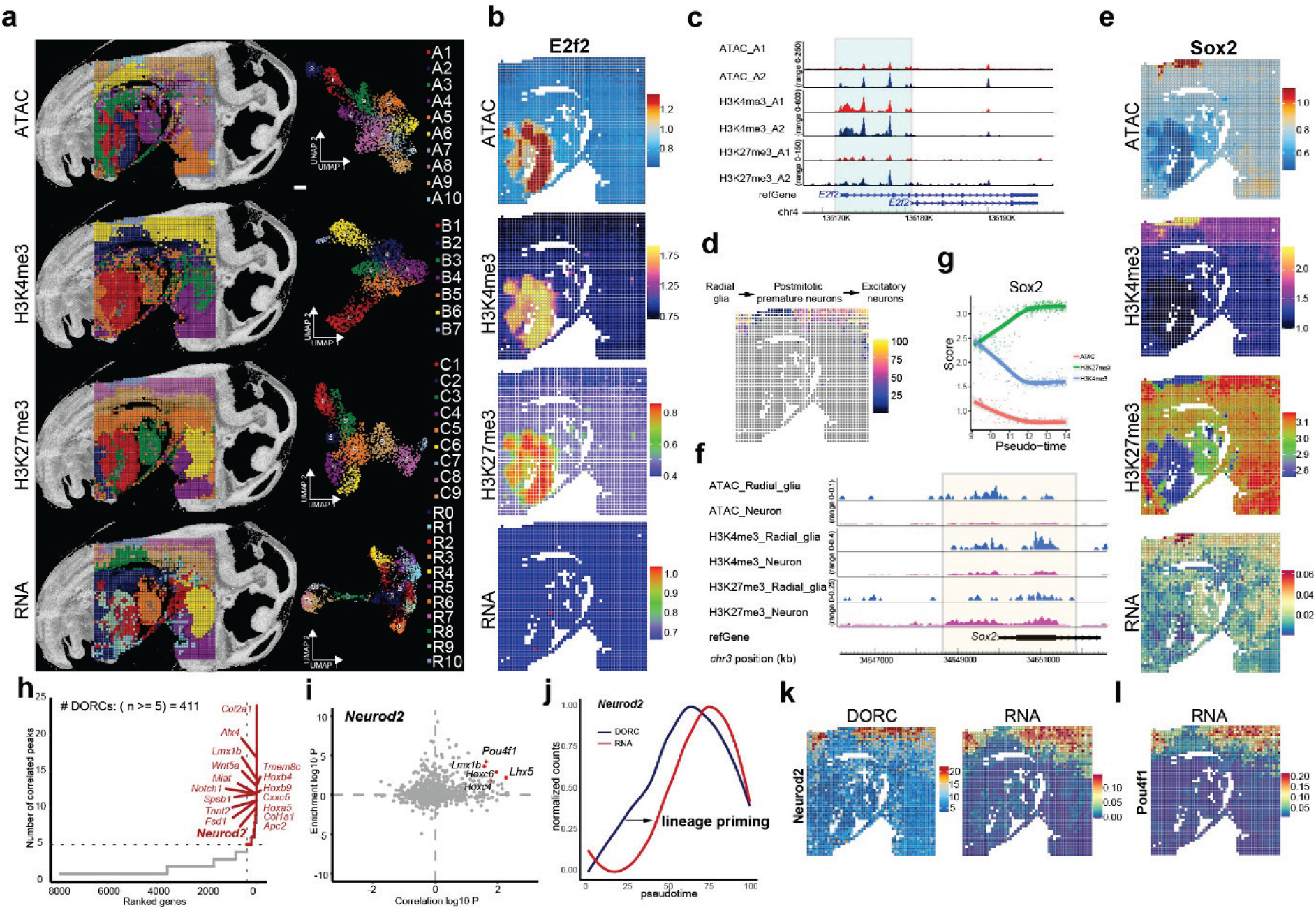
Spatial co-profiling of ATAC, RNA, H3K4me3, and H3K27me3 in mouse embryos. Sample: E13_50_μm_3. **a,** Spatial distribution and UMAP embeddings from unsupervised clustering analysis of four different modalities—ATAC-seq, RNA-seq, H3K4me3, and H3K27me3—profiling in E13 mouse embryos at a 50 μm pixel resolution. Distinct chromatin states and transcriptional landscapes in various embryonic regions, with clusters identified and visualized in anatomical context. **b,** Spatial mapping of *E2f2* gene with ATAC, RNA, H3K4me3 and H3K27me3 marks. **c,** Genome browser tracks of the *E2f2* gene showing chromatin accessibility (ATAC-seq), histone modifications (H3K4me3, H3K27me3), and RNA expression in liver clusters A1 and A2, as defined by ATAC-seq clustering. **d,** Integration of spatial ATAC-seq data with scRNA-seq data^14^ from E13.5 mouse embryos, followed by pseudotime analysis. The pseudotime trajectory from radial glia to postmitotic premature neurons and excitatory neurons is plotted in spatial coordinates, showing the dynamic chromatin landscape and transcriptional changes during neuronal differentiation. **e,** Spatial mapping of the *Sox2* gene across ATAC-seq, RNA-seq, H3K4me3, and H3K27me3 modalities, illustrating the multi-modal regulatory context of *Sox2* in the developing brain. **f,** Genome browser tracks of *Sox2* gene in ATAC, H3K4me3, and H3K27me3 modalities. The selected cell types are radial glia and postmitotic premature neurons. **g,** Scatter plot showing the dynamics of *Sox2* ATAC, H3K4me3, and H3K27me3 signals across pseudotime as determined in (**d**). The scaled scores reveal the temporal regulation of chromatin accessibility and histone modifications at the *Sox2* locus during neuronal differentiation. **h,** Spatial ATAC data and RNA data are used for domains of regulatory chromatin (DORCs) analysis with FigR^30^ package. The plot highlights the top-hit genes based on the number of significant gene-peak correlations across all cell types, emphasizing key regulatory elements in the embryonic genome. **i,** Identification of candidate transcription factor regulators of *Neurod2* using DORC analysis. Highlighted points represent top-hit transcription factors, indicating their potential regulatory influence on *Neurod2* expression during development. **j,** Comparison of chromatin (DORC) dynamics versus gene expression (RNA-seq) for *Neurod2*. This analysis illustrates the temporal relationship between chromatin state changes and transcriptional activation during lineage priming in neurodevelopment. **k,** Spatial patterns of DORCs *Neurod2* and its gene expression. **l,** Spatial gene expression of the transcription factor *Pou4f1*, showing its distribution in mouse embryonic neuronal development. Scale bar: 500 μm.

To further leverage the multimodal datasets, we conducted WNN analysis to integrate all trimodal and quadrimodal matrices (Extended Data Fig. 8). This approach enhanced the clustering identified by individual modality and revealed novel clusters that were not detectable with any single modality alone (Fig. 2a and Extended Data Fig. 8). For instance, the craniofacial region exhibited additional subclusters when analyzed through tri- or quadrimodal integration. Similarly, the heart region was further divided into two distinct subclusters through the integration of ATAC/H3K27me3/RNA or ATAC/H3K4me3/RNA modalities (Extended Data Fig. 8).

Recently, the co-profiling of chromatin accessibility and gene expression offers significant insights into the regulatory mechanisms of gene expression and cellular function^4,23^. However, there are situations that two modalities are not consistently correlated^4^, which could potentially be elucidated by considering additional epigenomic information. For example, *E2f1-3* genes were lowly expressed during fetal liver development^24,14^ (Extended Data Fig. 9a). despite, high chromatin accessibility was observed in the liver region (Fig. 2b and Extended Data Fig. 9b). This discrepancy could be explained by the co-measured H3K27me3 signals, which were also enriched at the promoter regions of *E2f* genes (Fig. 2c and Extended data Fig. 9c-d), indicating bivalency of *E2f* promoter in fetal liver.

We then annotated cell identities in each pixel by integrating the ATAC/H3K4me3 data with scRNA-seq mouse embryo dataset^14^, respectively. The spatial tissue pixels derived from both the ATAC data and H3K4me3 data were conformed well with the clusters of single-cell transcriptome (Extended Data Fig. 10a and b). We noted that chondrocytes and osteoblasts cluster A5 and B4), excitatory neurons (cluster A9 and B6), as well as radial glia (cluster A10 and B7), exhibited enrichment in the same spatial regions identified by both ATAC and H3K4me3 data. Remarkably, the ATAC data exhibited a greater abundance of postmitotic premature neurons compared to the H3K4me3 data, suggesting potential variations in the chromatin states of these adjacent neuron clusters.

To explore the spatiotemporal relationship between gene expression, chromatin accessibility, and histone modifications, we studied the developmental trajectory from radial glia to differentiated neurons^25^. A radial glia niche present in dorsal spinal cord could be revealed by all four modalities: cluster A10 of ATAC data, cluster B7 of H3K4me3 data, cluster C7 of H3K27me3 data, and cluster R10 of RNA data (Fig. 2a, Extended Data Fig. 10b). Pseudotime analysis^26^ was then conducted using the ATAC data, allowing for the visualization of the developmental trajectory on the tissue map (Fig. 2d). Several marker genes were identified and showed dynamic changes along this trajectory. For instance, *Sox2*, a master regulator of nervous system development and neuronal progenitors^27^, exhibited elevated chromatin accessibility and H3K4me3 within the radial glia whereas the levels of repressive H3K27me3 were low (Fig. 2e-g). Furthermore, spatial RNA data revealed region-specific gene expression of *Sox2* within the radial glia cluster. During the transition to postmitotic premature neurons and excitatory neurons, we observed a significant decrease in *Sox2* gene expression, along with the inaccessible chromatin, reduced H3K4me3 enrichment, and increased levels of H3K27me3. On the other hand, genes involved in neuronal development^28^ and synaptic transmission^29^, such as *Ank3* and *Gria2*, showed increased gene expression, along with accessible chromatin, consistent H3K4me3 enrichment, and low H3K27me3 enrichment at their gene loci (Extended Data Fig. 10c-e). We further analyzed Geno Ontology (GO) with spatial RNA data from radial glia and differentiated neuron clusters, and the results agreed with the anatomical annotation (Extended Data Fig. 10f-g).

During development, gene expression programs are orchestrated by a complex interplay between cis-regulatory elements and trans-acting factors, which together shape gene regulatory networks (GRNs). We integrated our multi-modal data for GRNs analysis using the FigR framework^30^,which links distal cis-regulatory elements with their target genes, facilitating the inference of GRNs and the identification of candidate transcription factor (TF) regulators that drive these networks. Analysis of co-profiled spatial ATAC-seq and RNA-seq datasets identified 411 lineage-determining genes marked as distinct domains of regulatory chromatin (DORCs)^31^ (Fig. 2h; Supplementary table 8). These DORCs were characterized by a high density of peak-gene associations and were significantly enriched for genes that play crucial roles in lineage determination and various developmental processes as confirmed by gene ontology (Extended data Fig. 11a). Among these, *Neurod2* stands out as a critical gene known for its pivotal role in guiding the differentiation of neural progenitor cells into mature neurons^32^. The spatial distribution of *Neurod2* showed high DORC accessibility and gene expression within clusters of postmitotic premature neurons and excitatory neurons (Fig. 2k), and changes in DORC accessibility of *Neurod2* preceded that of its gene expression along the differentiation trajectory due to the lineage-priming (Fig. 2j). We then calculated the enrichment of transcription factor motifs within the *Neurod2* DORC, to deduce potential TF activators (Fig. 2i). We identified *Pou4f1*, *Lhx5*, and *Lmx1b* as prominent transcriptional activators, whose involvement in dorsal spinal cord development have been described previously^33^. Their gene expression patterns were visualized across different tissue regions, demonstrating elevated expression levels specifically within these differentiated neurons (Fig. 2l and Extended Data Fig. 11b).

We further analyzed the GRN that is related to neurogenesis, and we identified that *Neurod2* could directly control *Nfib* expression (Extended Data Fig. 11c). Additionally, *Neurod2* and *Nfib* could co-regulate a set of genes, including *Sec14l1*, *Ap2a1*, and *Lingo1*, that were enriched in intermediate-stage neurons (Extended Data Fig. 11d). Collectively, our approach, offered a powerful tool to elucidate the regulatory mechanisms underlying development.

### Spatial co-profiling of protein expression, transcriptome, and histone modifications at near single cell resolution

H3K4me3 and H3K27me3 are two histone modifications with opposing roles in gene regulation. H3K4me3 is typically associated with active gene transcription, marking promoters of genes that are being expressed. In contrast, H3K27me3 is linked to gene repression, marking regions of the genome where gene expression is silenced. During development, the chromatin state where both gene-activating H3K4me3 and gene-repressing H3K27me3 marks co-occur at the promoters of developmental genes is known as bivalent chromatin^34^. This state involves regions marked simultaneously by these opposing histone modifications, keeping genes in a poised condition for rapid activation or repression. To date, the direct analysis of bivalent chromatin state and its effect on downstream gene and/or protein expression from the same sample at the genome scale and cellular level is lacking. We next performed the co-profiling of H3K27me3/H3K4me3, gene expression, and a panel of 7 cell surface proteins from the E13 hindbrain at near single cell resolution (E13_20_µm, Supplementary Table 7). We obtained a median of 1,510 (H3K27me3) and 897 (H3K4me3) unique fragments per pixel (Extended Data Fig. 12a-b) and observed the matched TSS enrichment scores of each modality (Extended Data Fig. 12c). For the RNA portion, total 22,165 genes were detected with an average of 1,258 genes and 1,999 UMIs per pixel (Extended Data Fig. 12b, 12e). To evaluate the impact of different pixel sizes on data quality, we compared samples E13_50_µm_3 and E13_20_µm, both derived from mouse embryonic day 13 tissue and sharing three modalities: H3K4me3, H3K27me3, and RNA. After downscaling to the same sequencing depth (50 million reads per sample), the 50-µm device showed higher unique fragment counts, gene counts, and UMIs than the 20-µm device, possibly due to capturing larger area and thus more nuclei per pixels, (Extended Data Fig. 12d-e).

Unsupervised clustering identified clusters with distinct spatial patterns: H3K27me3 clusters A1-A9, H3K4me3 clusters B1-B5, and RNA clusters R1-R12 (Fig. 3a), which agreed with tissue morphology (Fig. 3b). Each modality displayed similar clusters in the hindbrain but not in other regions, suggesting that H3K4me3 modifications may not be able to discriminate all cell types at this developmental stage. We then integrated spatial-RNA data with scRNA-seq data^14^ to assign cell types to each cluster (Fig. 3a-b, Extended Data Fig. 13a). Marker genes of spatial-RNA data identified major cell types, such as *Col1a* (osteoblasts), *Elavl2* (sensory neurons), *Hmga2* (epithelial cells), *Sox2/Pax3* (radial glia), and *Bcl11b* (postmitotic premature neurons).

**Fig. 3.**
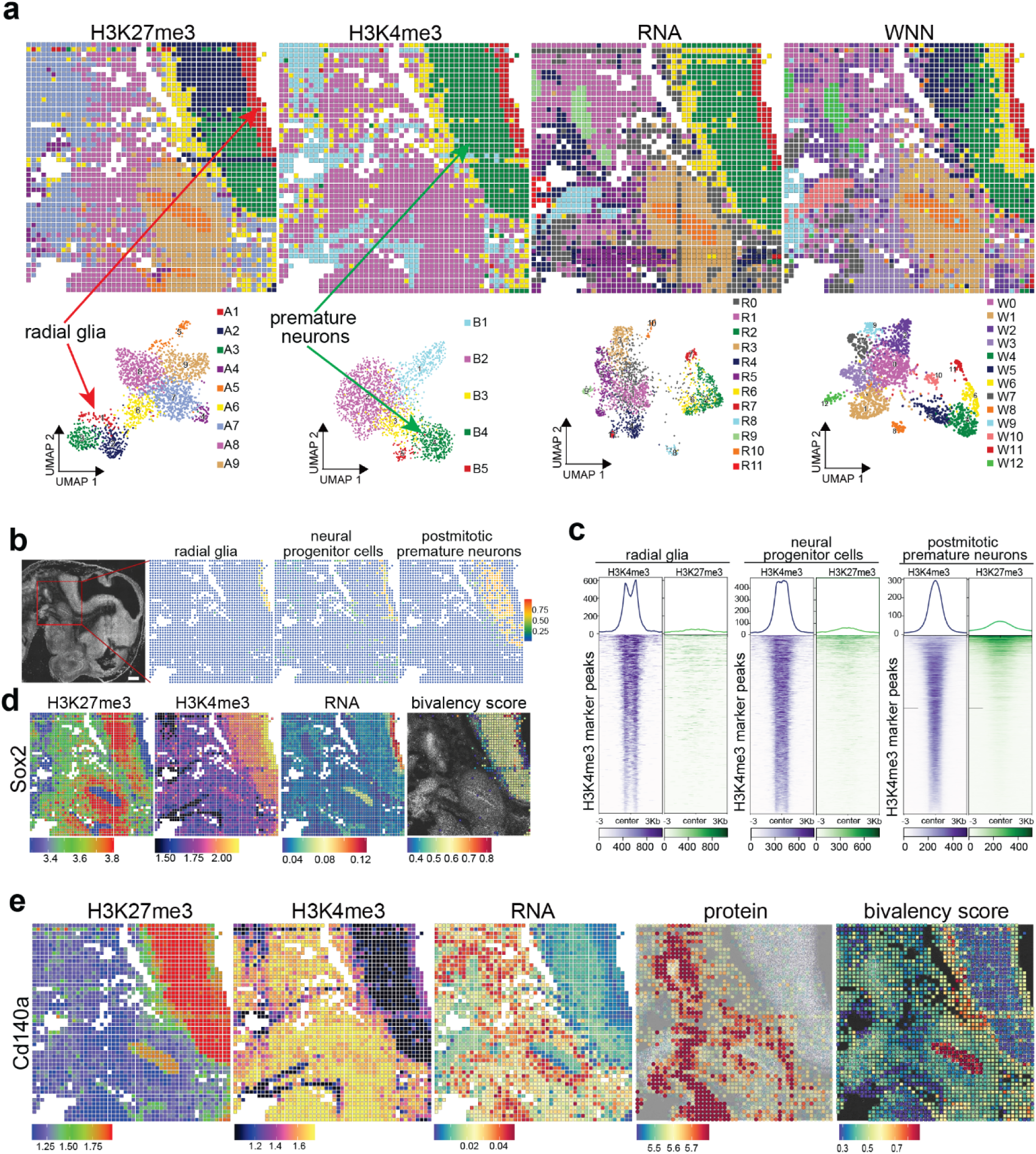
Spatial co-profiling of protein, RNA, H3K4me3, and H3K27me3 in mouse embryos. Sample: E13_20_μm. **a,** Spatial distribution and UMAP embeddings of unsupervised clustering analysis performed on each modality—H3K27me3, H3K4me3, RNA, and weighted-nearest neighbors (WNN) integration—at a 20 μm pixel resolution in E13 mouse embryos. Red arrow indicates the radial glia and green arrow indicates the premature neurons. **b,** Integration of spatial RNA data with scRNA-seq data^14^ from E13.5 mouse embryos enables high-resolution mapping of selected cell types, including radial glia, neural progenitor cells, and postmitotic premature neurons. This integration allows for the precise localization and characterization of these cell populations within the spatial context of the embryo. **c,** Deconvolution analysis of potential H3K4me3/H3K27me3 bivalency for clusters as determined in (**b**). **d,** Spatial mapping of the *Sox2* gene across RNA, H3K4me3, H3K27me3 modalities, and the calculated *Sox2* bivalency score. The bivalency score is derived from chromatin bivalency analysis, providing insight into the regulatory complexity of *Sox2* expression during neurodevelopment. The spatial distribution of *Sox2* bivalency highlights regions where the gene may be poised for activation or repression, depending on developmental cues. The bivalency score is calculated by chromatin bivalency analysis and described in **Methods**. **e,** Spatial patterns of the *Cd140a* gene, visualized across protein levels (using antibody-derived DNA tags), RNA expression, H3K4me3, H3K27me3, and the *Cd140a* bivalency score. This multi-modal spatial profiling reveals the complex regulatory environment of *Cd140a* and its role in embryonic development. The bivalency score provides additional context for understanding the interplay between chromatin state and gene expression in regulating *Cd140a*. Scale bar: 500 μm.

In the hindbrain region, the co-existence of various neuron types at different lineage stages makes it possible to explore the spatial temporal relationship between H3K4me3/H3K27me3 and gene/protein expression. We observed radial glia and postmitotic premature neurons were enriched in the similar clusters in the H3K27me3 (cluster A1-3) and H3K4me3 (cluster B4-5) (Fig. 3a). Moreover, the neural progenitor cells, derived from radial glia and endowed with self-renewal abilities to generate diverse neural cell types, could only be revealed by integrative analysis (Fig. 3a). To infer the dynamic change of potential H3K4me3/H3K27me3 bivalency during the transition of radial glia to differentiated neurons, we identified all active promoters specific to these neural cell types (Fig. 3b) and plotted the signals of H3K4me3 and H3K27me3 (Fig. 3c). Compared with neural progenitor cells and postmitotic premature neurons, radial glia showed the lowest enrichment of H3K27me3 signals in the promoter region that is defined by H3K4me3, reflecting lower level of bivalency or less heterogeneity of radial glia comparing to differentiating neurons.

To gain deeper insights into chromatin bivalency^35^ at cell type-specific gene loci, bivalency scores were used^36^,which provided a quantitative measure of the extent and intensity of bivalent chromatin domains at the level of individual cells or cell populations. By examining loci that function as markers for specific cell types, we were able to evaluate the dynamic interplay between gene-activating and gene-repressing histone modifications, particularly H3K4me3 and H3K27me3. For example, the bivalency score of the *Sox2* and *Pax3* genes, exhibited higher levels in postmitotic premature neurons compared to those in the radial glia cluster (Fig. 3d and Extended data Fig. 13b). It’s noteworthy that the precise regulation of gene expressions for *Sox2* and *Pax3* coincides with gradient changes, where there is an increase in H3K27me3 signals and a decrease in H3K4me3 signals. Another example is *Alx1*, where both its bivalency score and H3K4me3 signal decrease as cells differentiated (Extended Data Fig. 13b). Conversely, its H3K27me3 signal remained high, concurrent with the absence of *Alx1* gene expression.

In addition, we found that this four-modal profile was able to reveal the spatial patterns of cell surface proteins. For example, Cd140a protein was mainly detected within the non-neuronal region, which was concordant with its gene expression together with H3K4me3 and absence of H3K27me3 (Fig. 3e). In the epithelial cell cluster, the presence of a bivalent signal of H3K27me3/H3K4me3 at the *Cd140a* gene locus coincided with undetectable gene expression and absence of this surface protein. We subsequently visualized the expression of all seven individual proteins (Extended Data Fig. 14a-b). For example, the protein profiles of both Cd133 and B220 did not exhibit distinct spatial patterns, consistent with the spatial distribution observed in the Allen mouse brain *In Situ* Hybridization (ISH) datasets (Extended Data Fig. 14a-b). The spatial distribution of Cd90 proteins was assessed using antibodies specific to Thy-1.1 (Cd90.1) and Thy-1.2 (Cd90.2), which differ by a single amino acid^37^. As shown in the Extended Data Fig. 14c, Cd90.1 proteins exhibited a distinct distribution pattern in the hindbrain region. In contrast, Cd90.2 proteins demonstrated a broader distribution, with a noticeable presence in non-hindbrain regions. This differential expression underscores the importance of considering protein isoforms when assessing regional specificity during neurodevelopmental studies. In summary, spatial-Mux-seq enables the simultaneous measurement of modalities across two histone modifications, gene expression, and proteins from the same tissue section at nearly single-cell resolution.

### Multiplexed spatial mapping of mouse brain

Next, to evaluate the application of spatial-Mux-seq in different tissue types, we performed co-profiling of H3K27me3/H3K27ac and transcriptome of mouse postnatal day 21 hippocampus at near single cell resolution of 20 μm (P21_20_μm). A median of 3,571 (H3K27me3) and 1,249 (H3K27ac) unique fragments per pixel (Extended Data Fig. 15a-c) were obtained, and a total 23,090 genes were detected with an average of 1,499 genes and 2,848 UMIs per pixel (Extended Data Fig. 15b, 15e). We identified 11 H3K27me3 clusters (An), 10 H3K27ac clusters (Bn), and 9 RNA clusters (Rn) (Fig. 4a). These clusters agreed with the anatomical annotations in a hematoxylin and eosin (H&E)–stained adjacent tissue section (Fig. 4b). By integrating single-cell RNA-seq data^38^ from the mouse brain atlas with spatial RNA-seq data, we deconvoluted major cell types using RCTD^39^. Subsequently, we generated single-cell resolved cell-type maps across the mouse brain (Extended Data Fig. 15f). These maps revealed distinct spatial patterns that delineated various brain regions. For instance, both dentate gyrus granule neuroblasts and dentate gyrus granule neurons (DGGRC) were revealed in the dentate gyrus of the hippocampus, and CA excitatory neurons (TEGLU) were identified in the Cornu Ammonis region. Additionally, habenula cholinergic neurons (DECHO) and thalamus excitatory neurons (DEGLU) were found in thalamus with distinct spatial patterns.

**Fig. 4.**
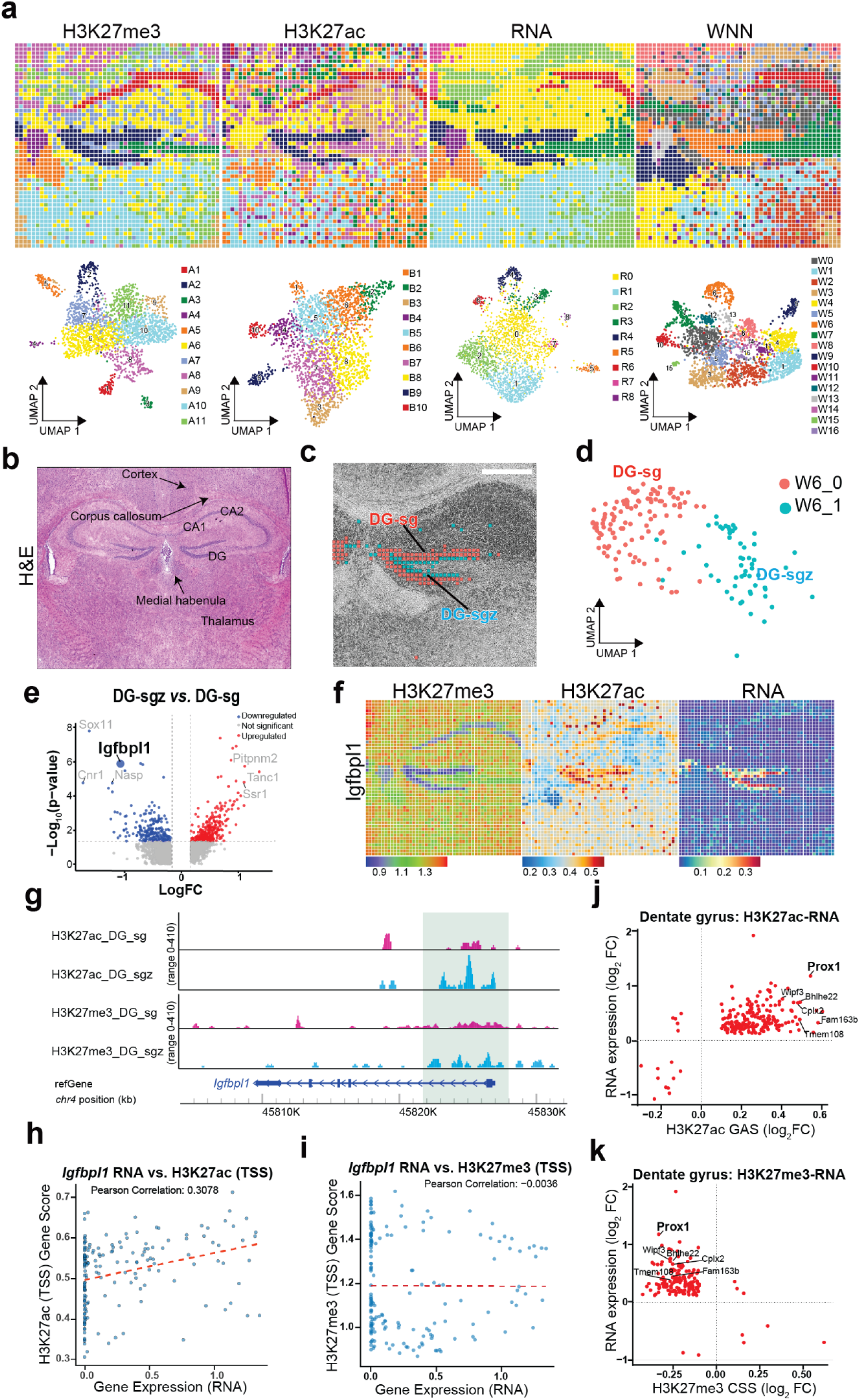
Spatial mapping of RNA, H3K27ac, and H3K27me3 in mouse juvenile brain. Sample: P21_20_μm. **a,** Spatial distribution and UMAP embeddings of unsupervised clustering analysis of H3K27me3, H3K27ac, RNA, and WNN with mouse juvenile brain (P21: 20 μm pixel size). Each modality provides a distinct perspective on the spatial organization of chromatin states and gene expression across the mouse brain region. **b,** Hematoxylin and Eosin (H&E) stained image of an adjacent tissue section from the juvenile mouse brain, providing anatomical context for the spatial molecular profiling data. **c,** Spatial mapping of two distinct hippocampal dentate gyrus subclusters: the dentate gyrus subgranular zone (DG-sgz) and the dentate gyrus granular cell layer (DG-sg). These subclusters represent specialized regions within the hippocampus, each with unique chromatin modifications and gene expression profiles. **d,** UMAP embeddings of the DG-sgz and DG-sg clusters, illustrating their distinct separation based on their molecular signatures. **e,** Differential expression of genes in DG-sgz clusters and DG-sg clusters. Volcano plot depicting the differentially expressed genes in DG-sgz clusters compared with DG-sg clusters*. P adj <0.05, logFC.threshold = 0.25*. **f,** Spatial mapping of the *Igfbpl1* gene, showing its expression across RNA, H3K27ac, and H3K27me3 modalities, and providing insight into its regulatory mechanisms within the hippocampus, particularly in the DG-sgz and DG-sg regions. **g,** Genome browser tracks for the *Igfbpl1* gene within the DG-sg and DG-sgz clusters, detailing the chromatin landscape at this locus. The tracks display the specific patterns of H3K27ac and H3K27me3 modifications, allowing for the comparison of active and repressive chromatin marks associated with *Igfbpl1* regulation in these hippocampal subclusters. **h-i,** Pearson correlation between *Igfbpl1* expression and histone mark H3K27ac (**h**) or H3K27me3 (**i**) gene scores. The gene scores are derived based on the gene model surrounding the transcription start site (TSS). (**g**) covering the DG-sg and DG-sgz clusters. **j,** Correlation of H3K27ac GAS and RNA gene expression. **k,** Correlation of H3K27me3 CSS and gene expression. Scale bar: 500 μm.

Building on these findings, we next examined the spatial patterns of specific markers to further distinguish cell types. As expected, we observed a robust enrichment of H3K27ac and elevated gene expression levels of *Mbp* specifically within the white matter of corpus callosum, whereas the H3K27me3 signal exhibited strongest intensity in the medial habenula region (Extended Data Fig. 16a). *Prox1* gene was highly expressed and was associated with strong enrichment of H3K27ac in the dentate gyrus of hippocampus. Remarkably *Prox1* was heavily marked by H3K27me3 specifically in the hippocampal CA region. Additional marker genes, such as *Scube1* and *Gria1*, exhibited specific H3K27me3 patterns in dentate gyrus or CA regions of hippocampus suggesting active involvement of H3K27me3 and polycomb repressive complex in the development of hippocampus in certain regions of the mouse brain (Extended Data Fig. 16a).

To further leverage the multimodal datasets, we performed WNN analysis by integrating the trimodal matrices. The integrative analysis effectively enhanced the clustering identified by each modality, and additionally captured novel clusters that could not be detected by any individual modality (Fig. 4a and Extended Data Fig. 16b). Within the thalamus region, further subdivision revealed three novel clusters: the stria medullaris (cluster W4), the central lateral nucleus of the thalamus (cluster W1), and the lateral dorsal nucleus of the thalamus (cluster W2). In adult mammals, radial glia-like cells generate granule cells from the dentate gyrus subgranular zone^40^. The maturation of granule cells occurs in the third postnatal week, which establishes a distinct granule cell identity^41^. To further reveal the diversity and molecular properties of mouse hippocampal progenitors, we subclustered the dentate gyrus granule cells and further identified two subclusters: dentate gyrus granule cell layer (DG-sg, cluster W6_0) and a thin layer of dentate gyrus granule subgranular zone (DG-sgz, cluster W6_1) (Fig. 4c-d). Differential gene expression analysis revealed that during the transition from DG-sgz to DG-sg, 228 genes were significantly downregulated, while 330 genes were significantly upregulated (*P adj < 0.05, logFC.threshold = 0.25*) (Fig.4e). For example, *Igfbpl1* expression was reduced in DG-sg relative to DG-sgz (Fig. 4f), whereas *Prox1* exhibited elevated expression in DG-sg compared to DG-sgz (Extended Data Fig. 16a). Upon analyzing their histone modifications along granular maturation, we noticed that the alteration in *Igfbpl1* expression coincided with a decrease in its H3K27ac signal without substantial increase in H3K27me3 (Fig. 4g-i), whereas the change observed in *Prox1* expression was associated with a decrease in H3K27me3 signal and an increase in H3K27ac signal (Extended Data Fig. 16c-e). In the hippocampal dentate gyrus, we observed a robust correlation between H3K27ac and gene expression and an anticorrelation between H3K27me3 and gene expression (Fig. 4j-k), including *Prox1*, *Wipf2*, and *Bhlhe22*, which exhibited significant enrichment with H3K27ac and minimal enrichment with H3K27me3, confirming the regulatory mechanism involving mutually exclusive H3K27me3/H3K27ac in gene expression regulation.

### Five-modal measurement of epigenome, transcription, and protein expression with spatial-Mux-seq

We finally sought to apply the spatial Mux-seq to enable simultaneous co-profiling of five modalities: chromatin accessibility, two histone modifications, transcriptome, and a large panel of cell surface proteins, in the same tissue section. By optimizing the sequential order of capturing different modalities, we were able to obtain ATAC/H3K27me3/H3K27ac libraries along with transcriptome and 122 oligo-tagged antibodies (Supplementary Table 7) from an adult mouse brain section (Extended Data Fig. 17a). Most of the oligo-tagged antibodies present in the commercial panel are immune markers and thus we specifically analyzed the mouse model of neuroinflammation-experimental autoimmune encephalomyelitis (EAE). EAE is an established and widely used model for multiple sclerosis that mimics many aspects of the human disease, including immune activation and infiltration into the central nervous system^42^. Using a 100 × 100 barcode scheme, the mapping area covered almost one hemisphere of the mouse brain in a coronal section. We obtained a median of 1,930 (ATAC), 1,433 (H3K27me3) and 405 (H3K27ac) unique fragments per pixel (Extended Data Fig. 17b-d), and a total 25,515 genes were detected with an average of 1,458 genes and 2,976 UMIs per pixel (Extended Data Fig. 17e). For the cell surface markers, we detected a median of 88 proteins and 728 protein UMIs per pixel (Extended Data Fig. 17e).

Unsupervised clustering was then performed for each modality separately, which identified 4 ATAC clusters (An), 11 H3K27me3 clusters (Bn), 8 H3K27ac clusters (Cn), 17 RNA clusters (Rn), and 7 protein clusters (Pn) (Extended Data Fig. 18a). Following this, we integrated the spatial ATAC, H3K27ac, and RNA data with single-cell RNA-seq reference data^38^ and identified major cell types (Extended Data Fig. 18b-c). For example, Medium Spiny Neurons (MSN1 and MSN2) were predominantly localized in the striatum, mature oligodendrocytes (MOL2) were found within the corpus callosum, and telencephalic glutamatergic neurons 8 (TEGLU8) were specifically distributed in the cortex.

To further validate these findings, we analyzed the spatial patterns of region-specific marker genes, confirming both the localization and functional relevance of the identified cell types (Extended Data Fig. 19). As expected, *Bcl11b* expression was predominantly observed in deep layer neurons and in the dorsal striatum, whereas it was repressed by H3K27me3 in superficial layers of cortex, corpus callosum and in the ventricular zone (Extended Data Fig. 19a). Interestingly, in contrast with the expression of *Bcl11b* mainly in dorsal striatum, H3K27ac was deposited on *Bcl11b* both in the dorsal and ventral striatum (Extended Data Fig. 19a). *Tbr1* expression, open chromatin and H3K27ac signal were mainly present in the cortex with anticorrelated H3K27me3 deposition (Extended Data Fig. 19a). *Dlx1* expression was detected in the lateral ventricle region, with more broad deposition of H3K27ac and chromatin opening also in the neighboring regions. Although we did not detect *Dlx1* expression in the striatum it was also not repressed by H3K27me3 there, whereas *Dlx1* repression by H3K27me3 occurred in the cortex (Extended Data Fig. 19a).

While the alignment of our integrated datasets with single-cell RNA sequencing data revealed a high degree of consistency between different modalities, the multifaceted nature of gene regulation might have some intriguing inconsistencies to be further studied. Specifically, when comparing chromatin accessibility, histone modifications, RNA, and protein expression, notable differences in the spatial patterns emerged (Extended Data Fig. 20a). In the corpus callosum, for example, the spatial patterns of Cd140a protein, RNA, ATAC-seq, and histone modifications revealed distinct variations. Cd140a protein expression exhibited a highly localized and defined pattern, contrasting with the more diffuse RNA signal. Interestingly, chromatin accessibility, as indicated by ATAC-seq, closely mirrored the protein expression pattern, suggesting that regions with accessible chromatin correlate with Cd140a protein localization. The histone modifications add another layer of complexity to this regulatory landscape. H3K27ac, typically associated with active enhancers, displayed a more widespread distribution, which did not directly correspond with the spatially well-defined expression of the Cd140a protein. In contrast, H3K27me3 exhibited a distinct and opposing spatial pattern, suggesting that certain *Cd140* isoforms might be epigenetically suppressed. Upon further analysis of individual Cd140 isoforms in the corpus callosum, we found that the longest *Cd140* isoform showed higher RNA expression, correlating with a lower H3K27me3 signal at its transcription start site, compared with other isoforms (Extended Data Fig. 20b). This suggests that the epigenetic landscape may selectively allow the transcription of certain isoforms while repressing others, highlighting the role of epigenetic mechanisms in precisely regulating gene expression.

## Discussion

The latest advances in spatial omics^4,7,43^, a rapidly evolving field, has enabled the investigation of complex biological systems with high-throughput quantifications of gene expression and epigenetic regulation within tissue context. However, gene and protein expression are regulated by different omics layers, such as DNA methylation^44^, chromatin remodeling^45^, histone modifications^46^, and genome architecture^47^. Despite recent single-cell technologies in trimodal measurements of RNA+ATAC+proteins^48,49^, H3K27me3+H3K27ac+protein^12^, or ATAC+H3K27me3+H3K27ac^9^, current spatial methods are limited to map two modalities at a time (such as ATAC+RNA^4,5^, CUT&Tag+RNA^4^, or protein+RNA^6–8^).

To overcome existing limitations in spatial multi-omics, we developed a novel technology, spatial-Mux-seq, that can simultaneously profile multiple histone modifications, chromatin accessibility, gene expression, and cell surface protein markers within the same tissue sections. This integrated approach provides a more comprehensive understanding of cellular states and regulatory mechanisms across spatial contexts. By co-profiling these modalities, spatial-Mux-seq enables the study of complex interplay between different regulatory layers, offering unprecedented insights into tissue architecture and function.

We rigorously benchmarked the spatial-Mux-seq datasets by comparing to previous methods, including spatial-CUT&Tag^11^, spatial-ATAC-RNA-seq, and spatial-CUT&Tag-RNA-seq^4^, evaluating them on key metrics such as the number of unique fragments, gene features, and UMIs. The results demonstrate that spatial-Mux-seq matches the performance of these techniques, confirming its capability to simultaneously profile multiple omics layers—histone modifications, chromatin accessibility, transcriptome, and proteins—without compromising the data quality from individual modality.

To demonstrate the versatility and accuracy of spatial-Mux-seq, we conducted four critical tests: 1. Histone modification co-profiling: We first validated the technology by co-profiling two mutually exclusive histone marks, H3K27me3 and H3K27ac. This test confirmed the accuracy and specificity of spatial-Mux-seq in capturing distinct epigenetic landscapes within the same tissue section. 2. Simultaneous profiling of four modalities: We simultaneously profiled two histone modifications (H3K27me3 and H3K4me3), transcriptome, and chromatin accessibility. This four-modality approach allowed us to track dynamic gene regulation from multi-layered epigenetic changes to gene expression, particularly during neural development in mice. 3. Integration of protein profiling: We extended spatial-Mux-seq to include a small panel of surface proteins, alongside mRNA and histone modifications (H3K4me3/H3K27me3), enabling simultaneous characterization of the epigenome, transcriptome, and proteome. This integration further demonstrates the broad applicability of spatial-Mux-seq in studying various aspects of gene regulation. 4. Comprehensive five-modality profiling: Finally, we applied spatial-Mux-seq to simultaneously measure chromatin accessibility, histone modifications (H3K27me3/H3K27ac), mRNA, and a large panel of 122 surface proteins within the same tissue section. The co-profiling of five modalities provides a more comprehensive view of cellular states and regulatory mechanisms, offering unparalleled insights into tissue biology.

By integrating multi-omics datasets, spatial-Mux-seq reveals a broader spectrum of cell types and uncovers connections between gene expression and various epigenetic changes. For instance, in the mouse hippocampus, our analysis of co-profiled H3K27me3, H3K27ac, and RNA data uncovered previously unrecognized roles for H3K27me3 in the maturation of dentate gyrus granular cells. Specifically, we observed increased transcriptional activity of the *Prox1* gene, essential for granule cell maturation, which was inversely correlated with H3K27me3 signals. This finding underscores the critical role of histone modifications in gene regulation and demonstrates the potential of spatial-Mux-seq to illuminate complex regulatory networks.

Despite these advancements, spatial-Mux-seq is currently limited to measuring two histone modifications at a time, primarily due to limitations in the restricted availability of nanobody-Tn5s^12^. Future improvements could overcome this limitation by developing additional nanobody-Tn5s from different species or by pre-conjugating primary antibodies with nanobody-Tn5s. Our study focuses on three critical histone marks: H3K27me3 (gene silencing), H3K4me3 (active promoters), and H3K27ac (active enhancers or promoters). While these marks are extensively used in epigenetic research for their significance in chromatin states and gene regulation, the exclusion of other histone marks may limit the scope of our conclusions. However, the selection was driven by antibody availability, reflecting technical constraints rather than a deliberate omission of other significant marks.

In conclusion, spatial-Mux-seq represents a significant advancement in spatial omics, offering a powerful tool for simultaneously assessing multiple regulatory layers within tissue context. By providing a more comprehensive understanding of complex biological systems and their underlying regulatory mechanisms, spatial-Mux-seq holds great promise for advancing our knowledge in fields such as developmental biology, disease research, and tissue engineering.

## Methods

### Preparation of tissue slides

Mouse C57 embryo sagittal frozen sections (MF-104-13-C57) were purchased from Zyagen. Tissue sections with 7-10 μm were collected on poly-L-lysine-coated glass slides. Juvenile mouse brain tissue (P21) was obtained from the C57BL/6 mice housed in the University of Pennsylvania Animal Care Facilities under pathogens-free conditions. All procedures used were pre-approved by the Institutional Animal Care and Use Committee. Juvenile mouse (P21) and adult mouse (5 months) were sacrificed by CO_2_, and brain was harvested and embedded in Tissue-Tek® O.C.T. compound (Sakura) and snap frozen using a mixture of dry ice and methylbutanol. The brains were coronally sectioned into 8 μm sections and collected on poly-L-lysine coated glass slides. The samples were stored at −80 °C until further use.

### Microfluidic device fabrication and assembly

The molds for polydimethylsiloxane (PDMS) microfluidic devices were fabricated using standard photolithography. The manufacturer’s guidelines were followed to spin-coat SU-8-negative photoresist (nos. SU-2025 and SU-2010, Microchem) onto a silicon wafer (no. C04004, WaferPro). The heights of the features were about 20 and 50 μm for 20- and 50-μm-wide devices, respectively. We mixed the curing and base agents in a 1:10 ratio and poured the mixture onto the molds. After degassing for 30 min the mixture was cured at 70 °C for 2 h. Solidified PDMS was extracted for further use. The fabrication and preparation of the PDMS device follow the published protocol^50^.

### Nanobody-Tn5 production and preparation of the Tn5 transposome

A detailed step-by-step protocol for purification of nanobody-Tn5 followed the published protocol^9^. Nanobody-Tn5 fusion proteins were loaded with barcoded oligonucleotides. The assemble process follows the published protocol^9^. Unloaded Tn5 transposase (C01070010) was purchased from Diagenode, and the transposome was assembled according to the manufacturer’s guidelines. The transposome was assembled by combination of Tn5MErev and Tn5ME-A or Tn5ME-B5/6/7. The oligo sequences used for transposome assembly were as follows:

Tn5MErev: 5′-/Phos/CTGTCTCTTATACACATCT-3′

Tn5ME-A: 5′-TCGTCGGCAGCGTCAGATGTGTATAAGAGACAG-3′

Tn5ME-B5 (wildtype Tn5): 5′-/Phos/CATCGGCGTACGACTTAGCCTAGATGTGTATAAGAGACAG-3′

Tn5ME-B6 (Mouse-nano-Tn5): 5’-/Phos/CATCGGCGTACGACTATAGAGAGATGTGTATAAGAGACAG-3′

Tn5ME-B7 (Rabbit-nano-Tn5): 5’-/Phos/CATCGGCGTACGACTCCTATCAGATGTGTATAAGAGACAG-3′

### DNA oligos, DNA barcode sequences and other key reagents

Lists of the DNA oligos that were used for sequencing library construction (N501, N7XX) and PCR (Supplementary Table 4), DNA barcode sequences (A1-100, B1-100) (Supplementary Table 5 and 6) and all other key reagents (Supplementary Table 7) are provided.

### Spatial co-profiling of ATAC, histone modifications, cell surface proteins, and gene expression

Frozen tissue slides were first thawed for 1 min at 37 °C. Tissue was fixed with formaldehyde (0.2%, with 0.05 U μl^-1^ RNase Inhibitor) for 5 min and quenched with 1.25 M glycine for a further 5 min. After fixation, tissue was washed twice with 1 ml of 1× DPBS-RI and cleaned with ddH_2_O. The sequential order for spatial multiple profiling is as follows: 1. ATAC-seq; 2.

Nanobody-based CUT & Tag; 3. Staining with cell surface markers 4. *In situ* reverse transcription; 5. Ligation of barcode A; 6. Ligation of barcode B; 7. Reverse crosslink; 8. gDNA and cDNA separation; 9. Library construction; 9. Library QC and sequencing. RNase Inhibitor (Enzymatics) was used in any buffers from step 1 to step 5 with a working concentration of 0.05 U μl^-1^. SUPERase•In™ RNase Inhibitor was used in Streptavidin C1 beads binding and washing processes.

1. **ATAC-seq:** Tissue section was permeabilized with lysis buffer (3 mM MgCl_2_, 0.01% Tween-20, 10 mM Tris-HCl pH 7.4, 0.01% NP40, 10 mM NaCl, 1% bovine serum albumin (BSA), 0.001% digitonin) for 15 min and washed twice with wash buffer (10 mM Tris-HCl pH 7.4, 10 mM NaCl, 3 mM MgCl_2_, 1% BSA, 0.1% Tween-20) for 5 min. Transposition mix (5 μl of home-made loaded Tn5 transposome, 33 μl of 1× DPBS, 50 μl of 2× Tagmentation buffer, 1 μl of 1% digitonin, 1 μl of 10% Tween-20, 10 μl of nuclease-free H_2_O) was added and incubated at 37 °C for 30 min. Next, 200 μl of 40 mM EDTA with 0.05 U μl^-1^ Enzymatic RNase inhibitor was added and incubated for 5 min at room temperature, to stop transposition.
2. **Nanobody-based CUT&Tag:** After ATAC process, the same tissue was washed twice with wash buffer (150 mM NaCl, 20 mM HEPES pH 7.5, one tablet of protease inhibitor cocktail, 0.5 mM Spermidine). The tissue section was then permeabilized with NP40-digitonin wash buffer (0.01% digitonin, 0.01% NP40 in wash buffer) for 5 min. The primary antibody (1:50 dilution with antibody buffer (0.001% BSA, 2 mM EDTA in NP40-digitonin wash buffer) was added and incubated at 4 °C overnight. A 1:100 dilution of nano-Tn5 adaptor complex mixture (rabbit-nano-Tn5/mouse-nano-Tn5) in 300-wash buffer (one tablet of Protease inhibitor cocktail, 300 mM NaCl, 0.5 mM Spermidine, 20 mM HEPES pH 7.5) was added and incubated at room temperature for 1 h, followed by a 5 min wash with 300-wash buffer. Tagmentation buffer (10 mM MgCl_2_ in 300-wash buffer) was added and incubated at 37 °C for 1 h. Next, 40 mM EDTA with 0.05 U μl^-1^ Enzymatic RNase inhibitor was added and incubated at room temperature for 5 min to stop the tagmentation process. The tissue was washed twice with 0.5× DPBS-RI for 5 min for further use.
3. **Staining with cell surface markers:** The tissue was washed twice with cell staining buffer and blocked with 1:20 mouse TruStain FcX™ in Cell Staining Buffer at 4°C for 15 min. Cell surface proteins are then detected with 1:400 oligonucleotide-labeled Antibody-Derived Tags (ADT) diluted in Cell Staining Buffer (1:400) at 4°C for 15 min, followed by a 5 min wash with Cell Staining Buffer. A 1:25 dilution of Fab Fragment (goat anti-mouse IgG) in Cell Staining Buffer was added and incubated at 4°C for 15 min. Discard reagent for further use.
4. ***In situ* reverse transcription:** The tissue was re-fixed with formaldehyde (2%, with 0.05 U μl^-1^ RNase Inhibitor) for 10 min and quenched with 1.25 M glycine for a further 5 min. The tissue was permeabilized with 0.5% Triton X-100 for 20min. The tissue was then washed twice with 0.5× DPBS-RI for 5 min. The tissue was then processed for mRNA detection and RT reaction, the following mixture was used: 12.5 μl of 5× RT buffer, 4.5 μl of RNase-free water, 0.4 μl of Enzymatic RNase inhibitor, 3.1 μl of 10 mM dNTP, 6.2 μl of Maxima H Minus Reverse Transcriptase, 25 μl of 0.5× PBS-RI and 10 μl of RT primer (100 μM). Tissues were incubated for 30 min at room temperature, then at 42°C for 90 min in a wet box. After the RT reaction, tissues were washed with 1× NEBuffer 3.1 containing 0.05 U μl^-1^ Enzymatic RNase inhibitor for 5 min.
5. **Ligation of barcode A::** Barcode A was pre-annealed with ligation linker 1, briefly, 10 μl of 100 μM ligation linker, 10 μl of 100 μM individual barcode A (A1-50 or A1-100) oligo and 20 μl of 2× annealing buffer (20 mM Tris pH 7.5–8.0, 100 mM NaCl, 2 mM EDTA) was mixed and reacted for annealing (95 °C for 5 min and cycling from 95 °C to 12 °C, 0.01 °C per cycle). For the first barcode (barcode A) *in situ* ligation, the PDMS chip A was covered to the region of interest (ROI). For alignment purposes, a 10× objective lens (BZ-X800 Series, Keyence) was used to take a brightfield image. The PDMS device and tissue slide were clamped tightly with a homemade acrylic clamp. Barcode A was pre-annealed with ligation linker 1, briefly, 10 μl of 100 μM ligation linker, 10 μl of 100 μM individual barcode A (A1-50, A1-100) oligo and 20 μl of 2× annealing buffer (20 mM Tris pH 7.5–8.0, 100 mM NaCl, 2 mM EDTA) was mixed and reacted for annealing. For each channel, 5 μl of ligation master mix containing individual barcode was loaded, it was prepared by mixing 2 μl of ligation mixture (27 μl of T4 DNA ligase buffer, 72.4 μl of RNase-free water, 5.4 μl of 5% Triton X-100, 11 μl of T4 DNA ligase), 2 μl of 1× NEBuffer 3.1 and 1 μl of each annealed DNA barcode A (A1−50 or A1-100, 25 μM). Vacuum was used to load the ligation master mix into 50 channels of the device, followed by incubation at 37 °C for 30 min in a wet box. The PDMS chip and clamp were removed after incubation and washed with 1× NEBuffer 3.1 containing 0.05U μl^-1^ Enzymatic RNase inhibitor for 5 min. Then the slide was washed with water and dried with compressed air.
6. **Ligation of barcode B:** Barcode B was pre-annealed with ligation linker 1, briefly, 10 μl of 100 μM ligation linker, 10 μl of 100 μM individual barcode B (B1-50 or B1-100) oligo and 20 μl of 2× annealing buffer (20 mM Tris pH 7.5–8.0, 100 mM NaCl, 2 mM EDTA) was mixed and reacted for annealing (95 °C for 5 min and cycling from 95 °C to 12 °C, 0.01 °C per cycle). For the second barcode (barcode B) *in situ* ligation, the PDMS chip B was covered to the ROI and a further brightfield image was taken with the 10× objective lens. An acrylic clamp was applied to clamp the PDMS, and the tissue slide together. Annealing of barcodes B (B1−50 or B1-100, 25 μM) and preparation of the ligation master mix were carried out as for barcodes A. The tissue was then incubated at 37 °C for 30 min in a wet box. After incubation, the PDMS chip and clamp were removed, and tissue was washed with 1× DPBS with Enzymatic RNase inhibitor for 5 min. The slide was then washed with water and dried with compressed air. A brightfield image covering each barcoding axis was then taken for further alignment.
7. **Reverse crosslink:** Lastly, the ROI on the tissue was digested with 100 μl of reverse crosslinking mixture (0.4 mg ml^-1^ proteinase K, 1 mM EDTA, 50 mM Tris-HCl pH 8.0, 200 mM NaCl, 1% SDS) at 58 °C for 2 h in a wet box. The lysate was then collected in a 0.2 ml tube and incubated at 60 °C overnight.
8. **gDNA and cDNA separation:** For gDNA and cDNA separation, the lysate was purified with Zymo DNA Clean & Concentrator-5 column and eluted with 100 μl of RNase-free water. 1× B&W buffer with 0.05% Tween-20 was used to wash 40 μl of Dynabeads™ MyOne™ Streptavidin C1 beads three times. Then, 100 μl of 2× B&W buffer with 2.5 μl of SUPERase•In™ inhibitor was used to resuspend the beads, which were mixed with the eluted DNA/cDNA mixture and allowed to bind the Biotinylated cDNA fragments at room temperature for 1 h with agitation.
9. **Library construction:** A magnetic rack was used to separate beads (containing cDNA/ADT) and supernatant (containing gDNA) in the eluent. The supernatant was collected and purified with with Zymo DNA Clean & Concentrator-5 again and eluted with 20 μl of RNase-free water for ATAC/nano-CUT&Tag library construction. 30ul of PCR mixture (25 μl of 2× NEBNext Master Mix, 2.5 μl of 10 μM indexed N7XX primer, 2.5 μl of 10 μM N501 PCR primer) was added to elute the gDNA. PCR reaction was first performed with the following program: 58 °C for 5 min, 72 °C for 5 min, 98 °C for 30 s and then cycling at 98 °C for 10 s, 60 °C for 30 s, 13 times. The final PCR product was purified by 1.3x SPRI beads (65 μl) and eluted in 20 μl of nuclease-free water. The separated beads were used for cDNA/ADT library construction. They were first washed twice with 400 μl of 1× B&W buffer with 0.05% Tween-20 containing 0.05 U/μl SUPERase•In™ RNase inhibitor and once with 10 mM Tris pH 8.0 containing 0.1% Tween-20 and 0.05 U μl^-^ SUPERase•In™ RNase inhibitor. The separated beads were then washed with 400 ul ddH_2_O. Streptavidin beads with bound cDNA/ADT molecules were resuspended in 200ul of TSO solution (22 μl of 10 mM deoxynucleotide triphosphate each, 44 μl of 5× Maxima RT buffer, 44 μl of 20% Ficoll PM-400 solution, 88 μl of RNase-free water, 5.5 μl of 100 uM template switch primer, 11 μl of Maxima H Minus Reverse Transcriptase, 5.5 μl of Enzymatic RNase Inhibitor) and were incubated at room temperature for 30 min and then at 42 °C for 90 min, with gentle shaking. After incubation, beads were washed once with 400 μl of 10 mM Tris and 0.1% Tween-20 and then with nuclease-free water. Washed beads were then resuspended in 220 ul of PCR solution (110 μl of 2× Kapa HiFi HotStart Master Mix, 8.8 μl of 10 μM PCR primer 1 and primer 2, 0.3 μl of 10 μM primer 3 (cite-seq), 92.4 μl of RNase-free water), then aliquoted 50ul beads mixture per PCR tube, and run on PCR thermocycling with the following program: 95 °C for 3 min and cycling at 98 °C for 20 s, 65 °C for 45 s and 72 °C for 3 min, 5 cycles. After the PCR reaction, beads were removed from the PCR product. 1× SYBR Green was added at a final concentration to the PCR product and run on a qPCR machine with the following thermocycling conditions: 95 °C for 3 min, cycling at 98 °C for 20 s, 65 °C for 20 s and 72 °C for 3 min, 15 times, followed by 5 min at 72 °C. The reaction was stopped once the qPCR curve signal began to plateau. The PCR product was then purified with 0.6x SPRI beads. The supernatant was saved for protein library (<200bp) and the separated SPRI beads were eluted in 20 μl of nuclease-free water for RNA library construction (>300bp). After all, a Nextera XT DNA Library Prep Kit was used for the RNA library generation. In brief, 1 ng of purified qPCR product was diluted in RNase-free water to a total volume of 5 μl, then 10 μl of Tagment DNA buffer and 5 μl of Amplicon Tagment mix were added and incubated at 55 °C for 5 min; 5 μl of NT buffer was then added to stop the tagmentation process, and incubated at room temperature for 5 min. 25 μl of PCR master solution (15 μl of PCR master mix, 1 μl of 10 μM N501 primer, 1 μl of 10 μM indexed N7XX primer, 8 μl of RNase-free water) was then added to the tagmentized DNA product and run with the following program: 95 °C for 30 s, cycling at 95 °C for 10 s, 55 °C for 30 s, 72 °C for 30 s and 72 °C for 5 min, for 12 cycles. The PCR product was purified with 0.6x SPRI beads to obtain the final RNA library. For protein library, the saved supernatant was purified with 1.4x SPRI beads and eluted in 20 μl of nuclease-free water. The eluted sample was repurified with 2.0x SPRI beads and finally eluted in 45 μl of nuclease-free water. PCR master solution (50 μl of 2× Kapa HiFi HotStart Master Mix, 2.5 μl of 10 μM P5 oligo (cite-seq), 2.5 μl of 10 μM indexed N7XX primer) was added to the eluted sample and performed the PCR reaction with the following program: 95 °C for 3 min, cycling at 95 °C for 20 s, 60 °C for 30 s, 72 °C for 20 s and 72 °C for 5 min, for 6 cycles. The PCR product was purified with 1.6x SPRI beads to obtain the protein library.
10. **Library QC and sequencing:** The Agilent D5000 Screentape was used to determine the size distribution and concentration of the library before sequencing. NGS was conducted on an Illumina NovaSeq 6000/NovaSeq X Plus sequencer (paired-end, 150-base-pair mode).

### Data preprocessing

For ATAC and CUT&Tag data, linkers 1 and 2 are used for targeted filtering of read 2, aligning utilizing BWA followed by sorting and indexing via Samtools to facilitate efficient data handling and retrieval. This reformatting process assigned genome sequences to the first read and incorporated barcodes A and B into the second read. We aligned these fastq files against mouse (GRCm38) reference genomes. The conversion produced tsv-like fragments files, enriched with spatial and genomic information through the integration of barcode pairs, facilitating comprehensive downstream analysis. For each modality, an ArchRProject was generated from the fragment file using ArchR v.1.0.2^51^ for downstream analysis. Peaks were called with pseudo-bulk bam files using MACS2 with parameters ‘--keep-dup=1 --llocal 100000 --min-length 1000 --max-gap 1000 --broad-cutoff=0.1’.

For RNA sequencing data, we refined read 2 to extract barcode A, barcode B, and the Unique Molecular Identifier (UMI). Using the Spatial Transcriptomics (ST) pipeline version 1.7.2, this processed data was mapped against the appropriate mouse (GRCm38) genome references. This step produced a gene matrix that captures both gene expression and spatial positioning data, encoded through the combination of barcodes A and B, enabling detailed spatial transcriptomic analysis. The gene matrix was then read into Seurat v.4.3.0^13^ as a Seurat object.

For cDNAs derived from ADTs, the raw FASTQ file of Read 2 was reformatted the same way as cDNAs from RNA. Using default settings of CITE-seq-Count 1.4.2^52^, we counted the ADT UMI numbers for each antibody in each spatial location. The protein expression matrix contains the spatial locations (barcode A × barcode B) of the proteins and protein expression levels.

### Data clustering and visualization

Firstly, we identified the location of pixels on tissue from the brightfield image using a custom python script (https://github.com/liranmao/Spatial_multi_omics).

For ATAC and CUT&Tag data, based on the ArchRProject, the normalization and dimension reduction were conducted using LSI and UMAP. Then we used the getGeneScore from ArchR package to get the GAS and the CSS scores. For spatial data visualization, to facilitate the mapping of data onto the original tissue, the gene score matrix derived from ArchR was imported into Seurat as a Seurat object. Then we ploted the spatial maps using SpatialPlot. The size of the pixels was adjusted for visualization by modifying the ‘pt.size.factor’ parameter within the Seurat package.

For RNA data, based on the Seurat object, we used the SCTtransform function for the data normalization and variance stabilization. Then the dimensionality reduction was done by RunPCA. We then constructed the nearest neighbor graph on the first 30 PCs by the function FindNeighbors. The clusters were identified with appropriate resolutions. Ultimately, we computed a UMAP embedding leveraging the initial 30 principal components using RunUMAP. And SpatialPlot was used for spatial plot visualization.

Protein data were normalized using the centered log ratio (CLR) transformation method in Seurat version 4.3.0. All heat maps were plotted using ggplot2. And SpatialPlot was used for spatial plot visualization. This was the same as ATAC and CUT&Tag data.

### Multi-omics integration

For our multi-omics data integration, we consolidated ATAC, CUT&Tag, and RNA datasets into a single Seurat object. The ATAC and CUT&Tag data integration utilized a 500bp peak matrix generated by addReproduciblePeakSet from ArchR, applying Macs2 for peak calling. RNA data integration was based on a log-normalized gene expression matrix. We applied Weighted Nearest Neighbors (WNN) analysis with FindMultiModalNeighbors for clustering, utilizing UMAP and spatial mapping for visualization. Subsequently, cell type clusters were refined through FindSubCluster within Seurat, based on the wsnn graph. This streamlined approach facilitated a precise analysis of cellular heterogeneity within the multi-omics dataset.

### Integrative data analysis and cell type identification

To delineate cell identities within each pixel, we employed the addGeneIntegrationMatrix function from ArchR, integrating ATAC/H3K4me3 data with a single-cell RNA-seq. To get a higher resolution cell type inference inside one pixel, we used robust cell type decomposition (RCTD)^39^ to decompose cell-type mixtures by leveraging cell type profiles learned from single-cell RNA-seq.

### Downstream analysis

For assessing the correlation of CSS/GAS and gene expression, we performed the analysis for certain identified cell type clusters, dentate gyrus specifically. Marker genes from the RNA dataset were identified using the FindMarkers function, applying the Wilcoxon rank sum test with a log_2_ fold change threshold of 0.10. We further filtered the RNA markers based on an adjusted P-value threshold of < 10^-5^. Similarly, for chromatin features, including gene activity score (GAS), and chromatin silencing score (CSS)), we employed the FindMarkers function with identical parameters to determine the marker genes. Gene Ontology (GO) analysis was conducted using enrichGO function from R package clusterProfiler v4.8.3^53^.

### Chromatin dynamics analysis

Pseudo-time analysis on RNA was performed using Slingshot v2.2.1. The trajectory analysis on ATAC was conducted employing addTrajectory function from ArchR. For chromatin bivalency analysis, we considered genes exhibiting high levels of both H3K4me3 and H3K27me3 as bivalent. For a certain gene, the H3K4me3 and H3K27me3 signal of each pixel were calculated by getGeneScore function from ArchR package, identifying the subset of signals that were within the gene window weighted the distance. The bivalency score was calculated as previously published method^36^.

### Gene regulation analysis

We used FigR v0.1.0^30^ to infer the transcriptional regulation by integrating ATAC and RNA data. The runGenePeakcorr function facilitated peak-gene association testing. Domains of regulatory chromatin (DORCs) were defined as genes with a relatively high number of significant peak-gene associations (n>=5). DORC accessibility scores were obtained using the getDORCScores function. To pinpoint potential transcription factors (TFs) regulating DORCs, the runFigGRN function was employed to identify TF binding motifs enriched within specific DORCs, indicating their potential role in driving DORC regulation.

## Code availability

The whole analysis pipeline and instructions for reproduction are available on Github (https://github.com/liranmao/Spatial_multi_omics).

## Data availability

Raw and processed data reported in this paper are deposited in the Gene Expression Omnibus (GEO) with accession code GSE263333. Resulting fastq files were aligned to the mouse reference genome (mm10). Published data for data quality comparison and integrative data analysis include: Mouse organogenesis cell atlas (MOCA) (https://oncoscape.v3.sttrcancer.org/atlas.gs.washington.edu.mouse.rna/downloads), ENCODE mouse embryo H3K27me3 and H3K27ac chip-seq datasets (13.5 days) (https://www.encodeproject.org/), mouse brain cell atlas (http://mousebrain.org/adolescent/downloads.html), and Allen Developing Mouse Brain Atlas (https://developingmouse.brain-map.org/).

## Acknowledgments

We acknowledge support from the Packard Fellowship for Science and Engineering (to Y.D.), John Q. Trojanowski Research Scholar Award from the Penn Institute on Aging (to Y.D.), and the US National Institutes of Health (grant numbers DP2AI177913 to Y.D.).

## Contributions

Methodology: P.G., M.B., and Y.D.; Experimental Investigation: P.G., Y.C., C.N.L., and A.C.; Data Analysis: P.G., L.M., M.L., and Y.D.; Original Draft: P.G., L.M., and Y.D. All authors reviewed, edited, and approved the manuscript.

## Competing interests

Y.D. and P.G. are inventors of a patent application related to this work. M.L. receives research funding from Biogen Inc. unrelated to the current manuscript and is a co-founder of OmicPath AI LLC. The other authors declare no competing financial interests.

---

**Supplementary Table 1.**
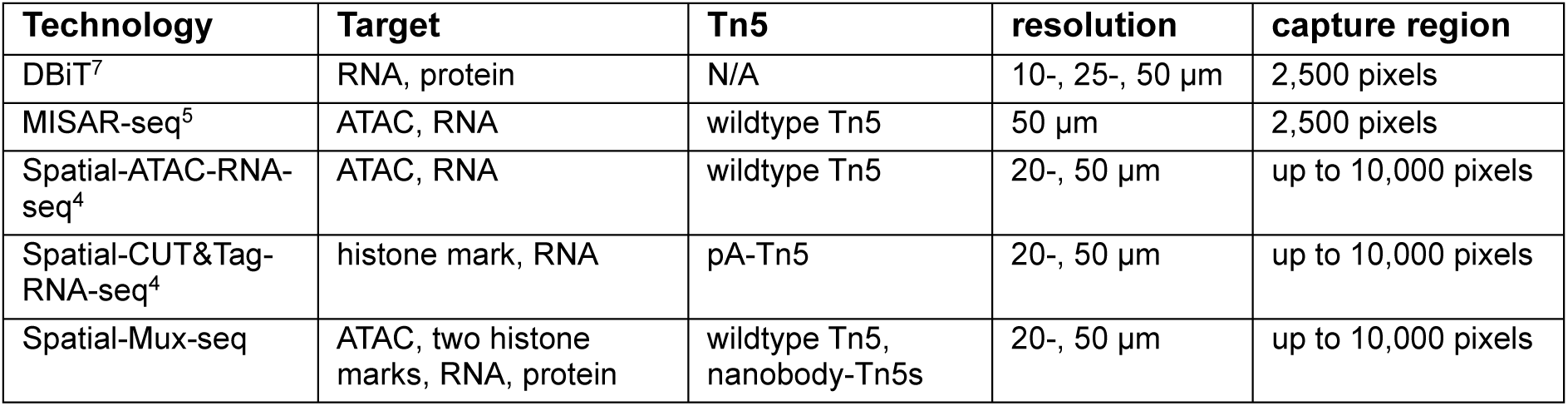
Comparison of spatial multi-omics methods utilizing microfluidic devices.

| Technology | Target | Tn5 | resolution | capture region |
| --- | --- | --- | --- | --- |
| DBiT<sup>7</sup> | RNA, protein | N/A | 10-, 25-, 50 µm | 2,500 pixels |
| MISAR-seq<sup>5</sup> | ATAC, RNA | wildtype Tn5 | 50 µm | 2,500 pixels |
| Spatial-ATAC-RNA-seq<sup>4</sup> | ATAC, RNA | wildtype Tn5 | 20-, 50 µm | up to 10,000 pixels |
| Spatial-CUT&Tag-RNA-seq<sup>4</sup> | histone mark, RNA | pA-Tn5 | 20-, 50 µm | up to 10,000 pixels |
| Spatial-Mux-seq | ATAC, two histone marks, RNA, protein | wildtype Tn5, nanobody-Tn5s | 20-, 50 µm | up to 10,000 pixels |

**Supplementary Table 2.** Summary of spatial multi-modalities profiling of all the samples.

Summary of spatial multi-modalities profiling of all the samples.
| Sample | tissue type | Spatial Resolution | Modalities |
| --- | --- | --- | --- |
| E13_50_µm_1 | mouse embryo day 13 | 50 µm, 2500 pixels | H3K27me3, H3K27ac |
| E13_50_µm_2 | mouse embryo day 13 | 50 µm, 2500 pixels | H3K27me3, H3K27ac |
| E13_50_µm_3 | mouse embryo day 13 | 50 µm, 2500 pixels | ATAC, H3K4me3, H3K27me3, RNA |
| E13_20_µm | mouse embryo day 13 | 50 µm, 2500 pixels | H3K27me3, H3K4me3, RNA, protein |
| P21_20_µm | mouse brain day 21 | 20 µm, 2500 pixels | H3K27me3, H3K27ac, RNA |
| 5M_20_µm | mouse brain 5 months | 20 µm, 10000 pixels | ATAC, H3K27me3, H3K27ac, RNA, protein |

**Supplementary Table 3.**
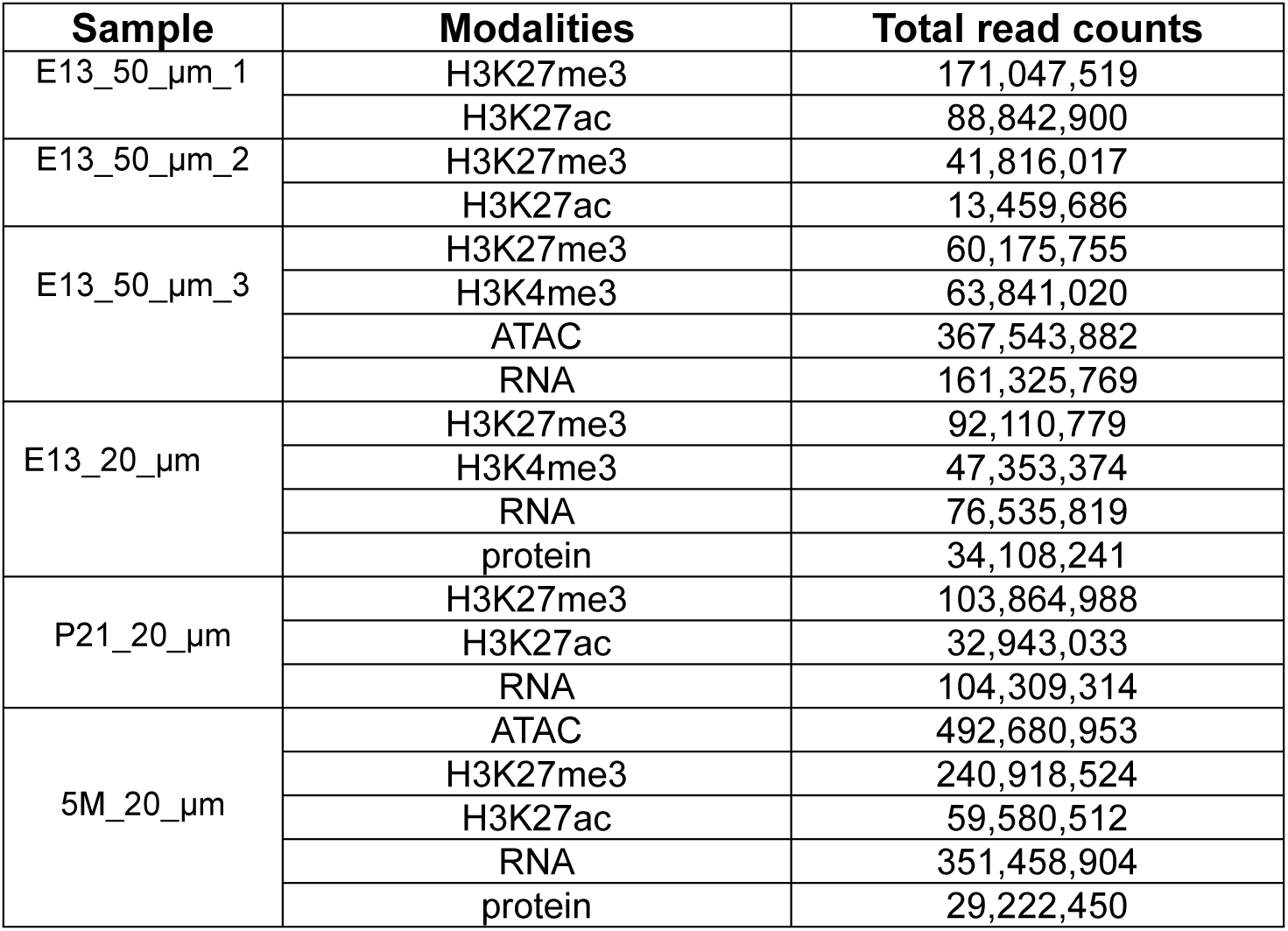
Summary of sequence depths for spatial-Mux-seq profiling of all the samples.

**Supplementary Table 4.**
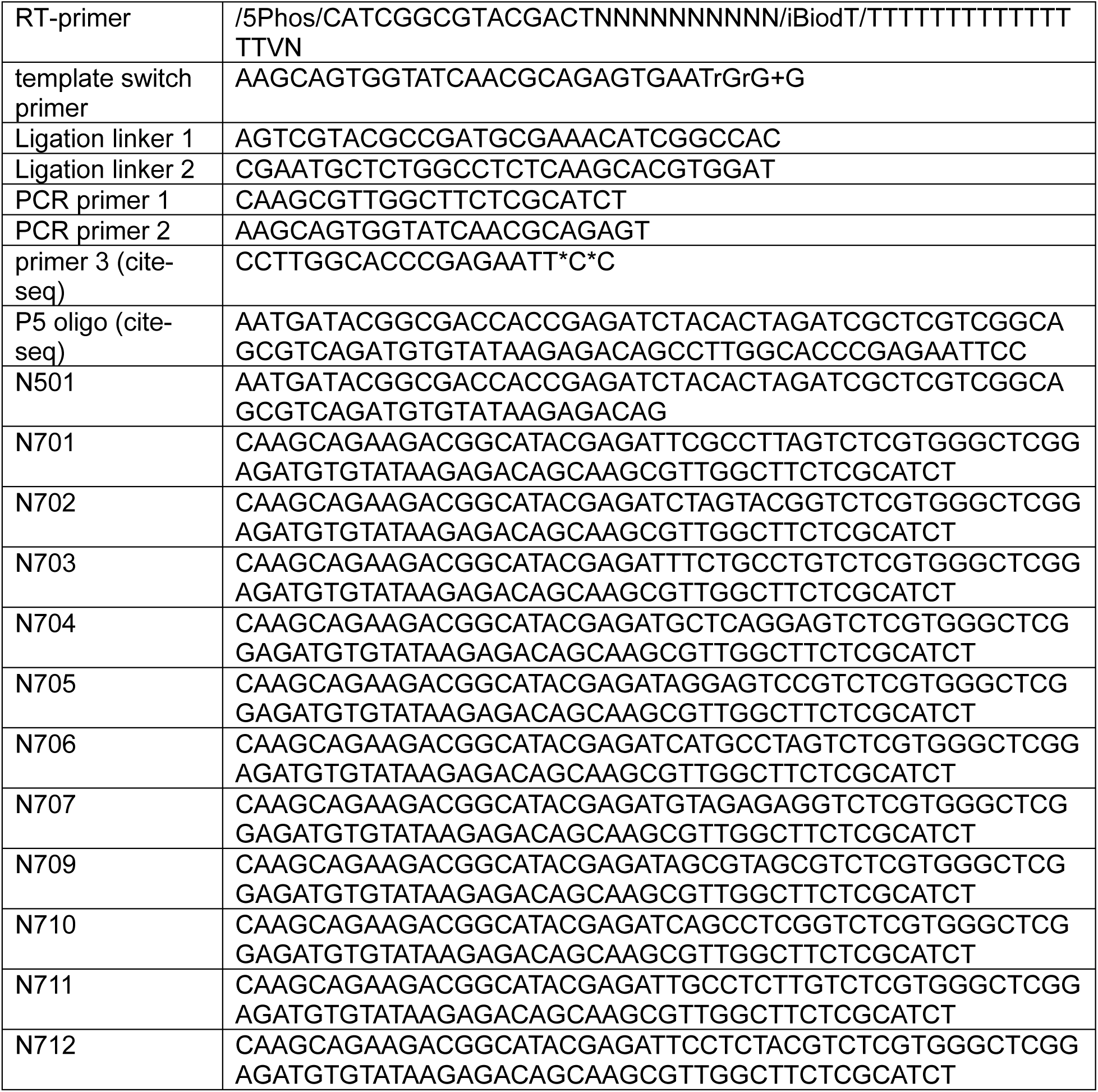
DNA oligos used for PCR and preparation of sequencing library.

**Supplementary Table 5.**
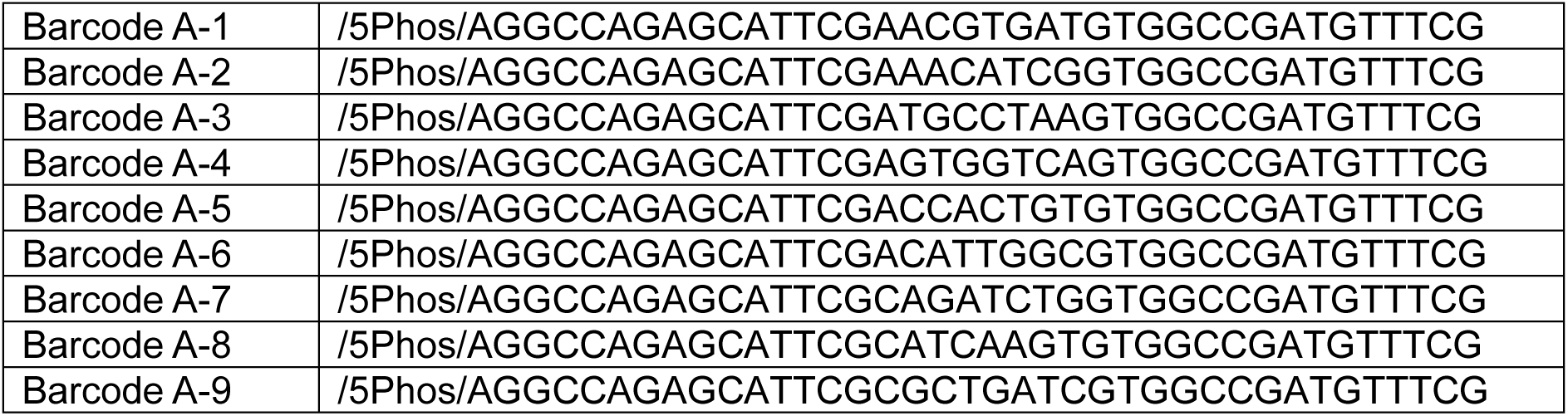
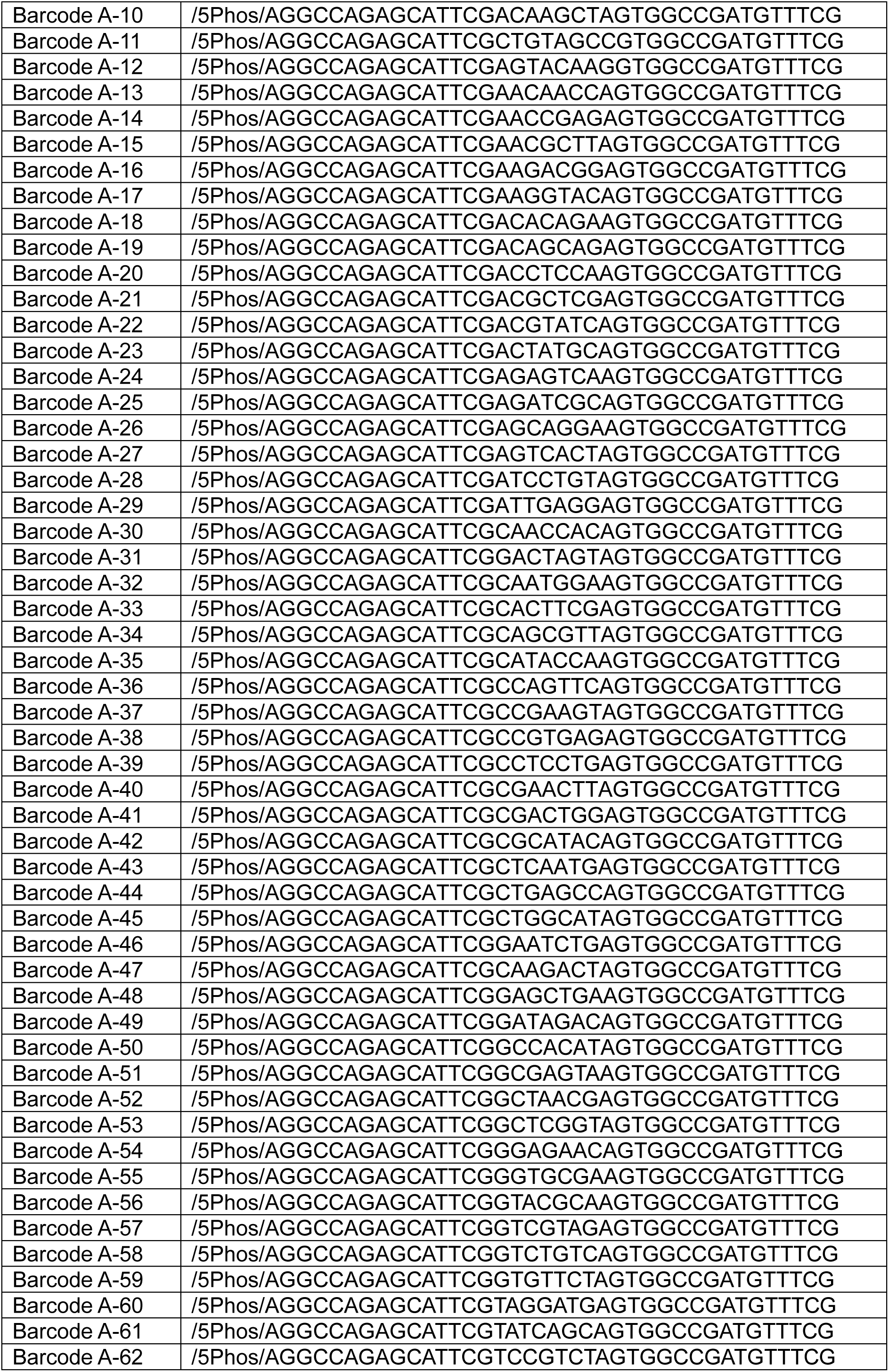
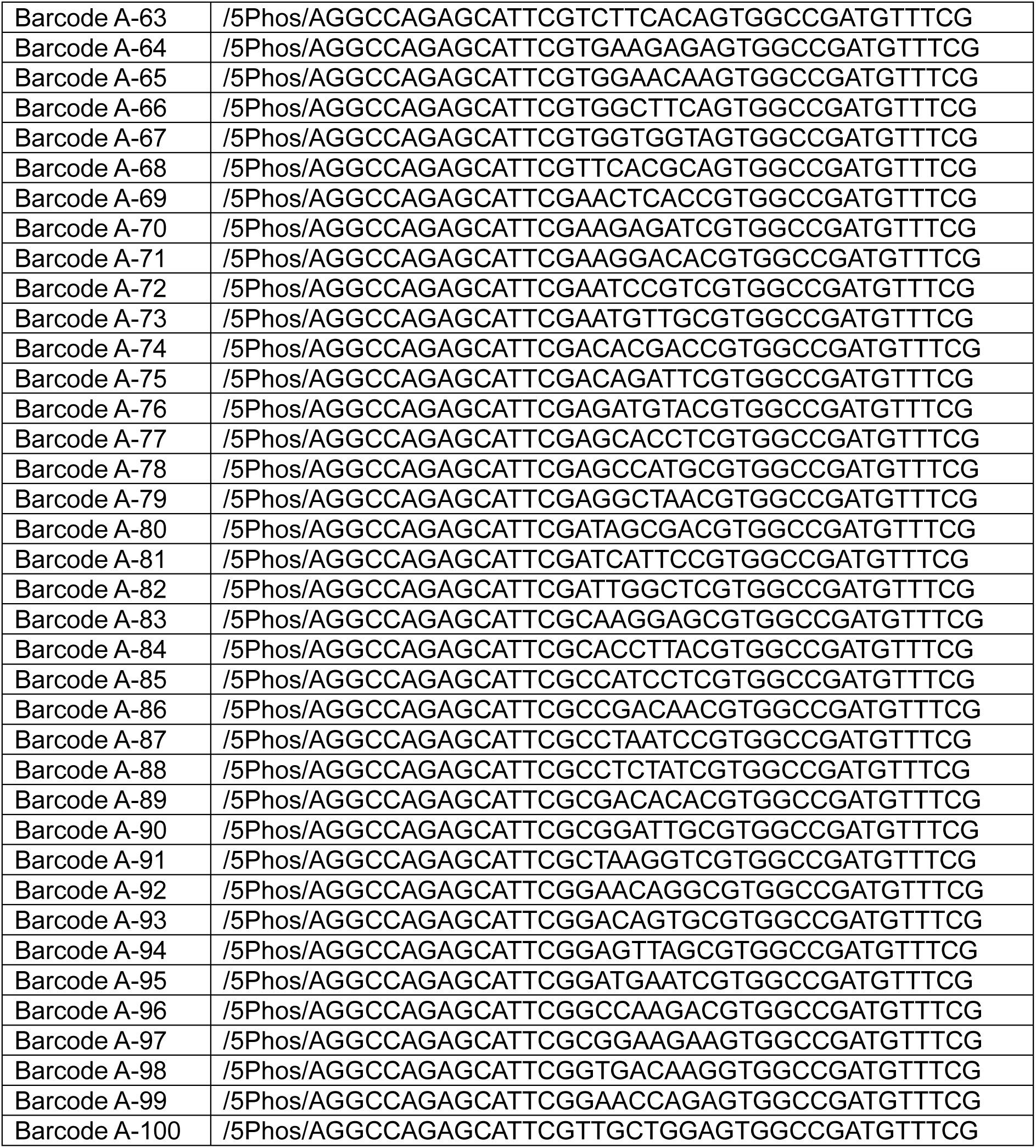
Barcode A Sequence.

**Supplementary Table 6.**
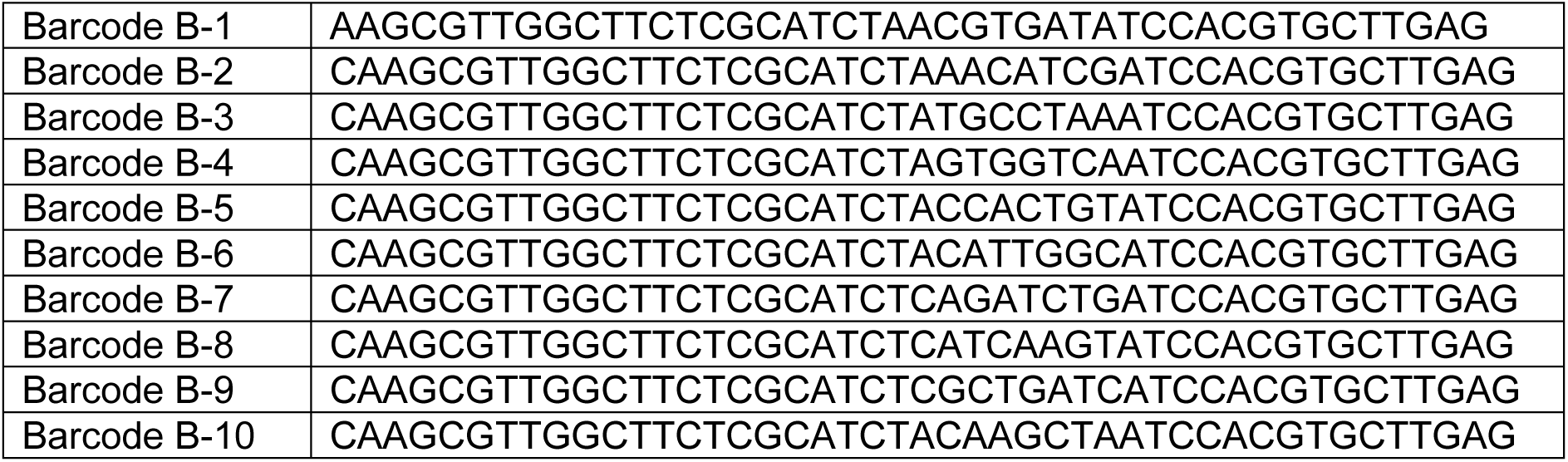
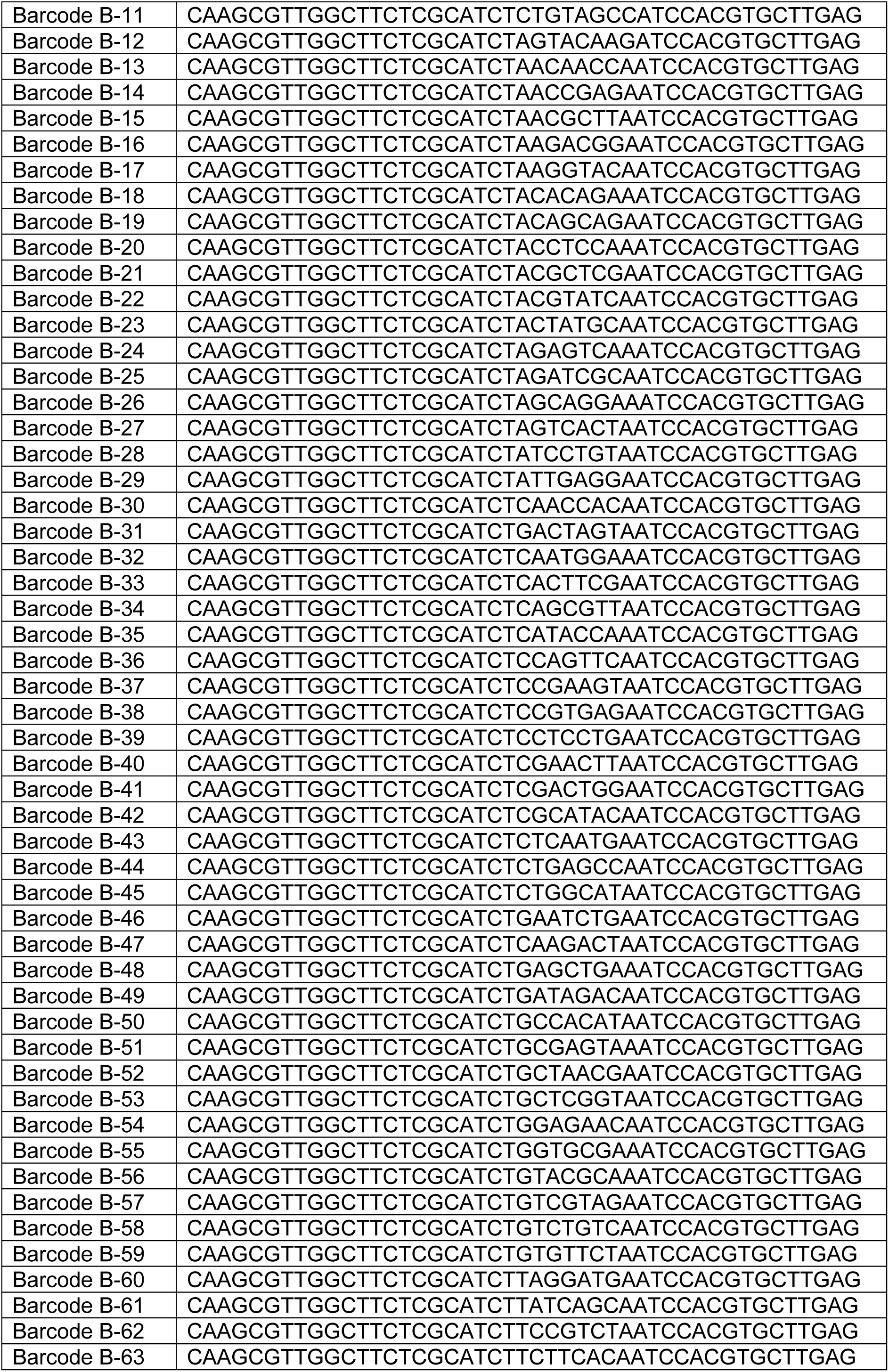
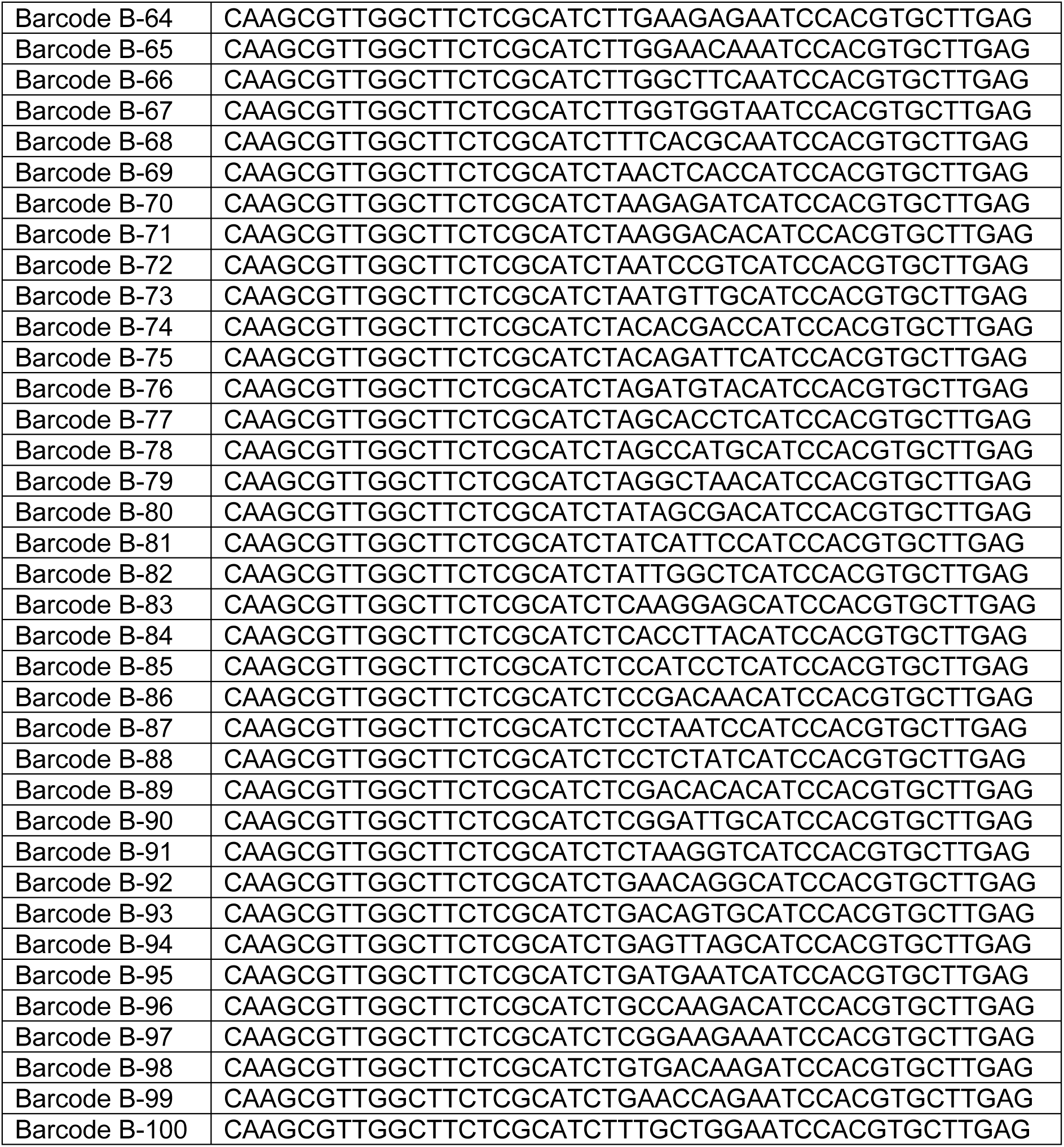
Barcode B Sequence.

**Supplementary Table 7.** Chemicals and reagents.

| Name | Catalog number | Vender |
| --- | --- | --- |
| Formaldehyde solution | PI28906 | Thermo Fisher Scientific |
| HEPES pH 7.5 | BBH-75-250 | Boston BioProducts |
| Glycine | 50046 | Sigma-Aldrich |
| NaCl | AM9760G | Thermo Fisher Scientific |
| Digitonin | G9441 | Promega |
| MgCl<sub>2</sub> | AM9530G | Thermo Fisher Scientific |
| Spermidine | S0266 | Sigma-Aldrich |
| EDTA-free Protease Inhibitor Cocktail | 11873580001 | Millipore Sigma |
| NP40 | 11332473001 | Sigma-Aldrich |
| EDTA Solution pH 8.0 | AB00502 | AmericanBio |
| Bovine Serum Albumin (BSA) | A8806 | Sigma-Aldrich |
| Anti-H3K27me3 antibody | Ab6002 | Abcam |
| Anti-H3K27ac antibody | 8173 | Cell Signaling Technology |
| Anti-H3K4me3 antibody | 9751 | Cell Signaling Technology |
| TotalSeq™-A Mouse Universal Cocktail | 199901 | Biolegend |
| Anti mouse CD4 | A0001 | Biolegend |
| Anti mouse CD3 | A0182 | Biolegend |
| Anti mouse CD34 | A0857 | Biolegend |
| Anti mouse CD140a | A0573 | Biolegend |
| Anti mouse CD133 | A1037 | Biolegend |
| Anti mouse CD90.1 | A0380 | Biolegend |
| Anti mouse CD90.2 | A0075 | Biolegend |
| Anti mouse B220 | A0103 | Biolegend |
| TruStain FcX™ PLUS (anti-mouse CD16/32) antibody | 156604 | Biolegend |
| Cell Staining Buffer | 420201 | Biolegend |
| Fab Fragment Goat Anti-Mouse IgG | 115-007-003 | Jackson ImmunoResearch |
| Triton X-100 | T8787 | Sigma-Aldrich |
| T4 DNA Ligase | M0202L | New England Biolabs |
| T4 DNA Ligase Reaction Buffer | B0202S | New England Biolabs |
| NEBuffer 3.1 | B7203S | New England Biolabs |
| DPBS | 14190144 | Thermo Fisher Scientific |
| Proteinase K | EO0491 | Thermo Fisher Scientific |
| SPRI beads | A63880 | Beckman Coulter |
| NEBNext High-Fidelity 2X PCR Master Mix | M0541L | New England Biolabs |
| SYBR Green I Nucleic Acid Gel Stain | S7563 | Thermo Fisher Scientific |
| DNA Clean & Concentrator-5 | D4014 | Zymo Research |
| Tn5 Transposase - unloaded | C01070010 | Diagenode |
| Tagmentation Buffer (2x) | C01019043 | Diagenode |
| Sodium dodecyl sulfate | 71736 | Sigma-Aldrich |
| Maxima H Minus Reverse Transcriptase (200 U/L) | EP0751 | Thermo Fisher Scientific |
| dNTP mix | R0192 | Thermo Fisher Scientific |
| SUPERase•In™ RNase Inhibitor | AM2694 | Thermo Fisher Scientific |
| Dynabeads™ MyOne™ Streptavidin C1 | 65001 | Thermo Fisher Scientific |
| RNase Inhibitor | Y9240L | Enzymatics |
| Kapa Hotstart HiFi ReadyMix | KK2601 | Kapa Biosystems |
| Nextera XT DNA Library Preparation Kit | FC-131-1024 | Illumina |

