## Extended Data Figures for "Multiplexed spatial mapping of chromatin features, transcriptome, and proteins in tissues"

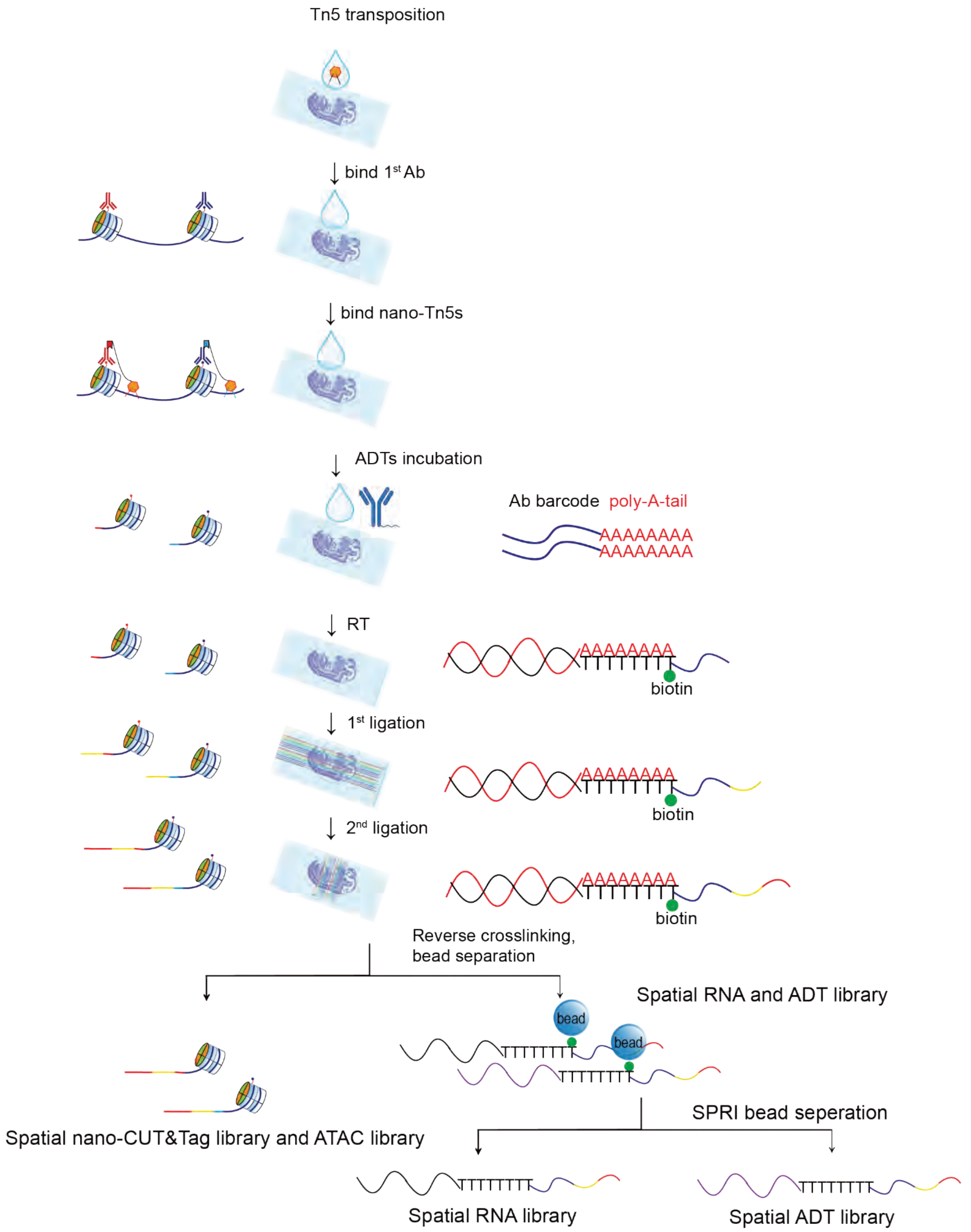


**Extended Data Fig. 1 | Workflow of spatial-Mux-seq.** Schematic workflow for spatial co-profiling of ATAC, two histone modifications, transcriptomes, and cell surface proteins: A tissue section was first incubated with wildtype Tn5. Two primary antibodies against different histone marks were then added, followed by incubation with two secondary nanobody-Tn5s. Next, a panel of ADTs is used to label cell surface proteins. *In situ* reverse transcription was then performed, followed by two rounds of DNA barcoding to create a mosaic of tissue pixels. Finally, gDNA and cDNA were collected and separated, and library construction was completed with PCR amplification.

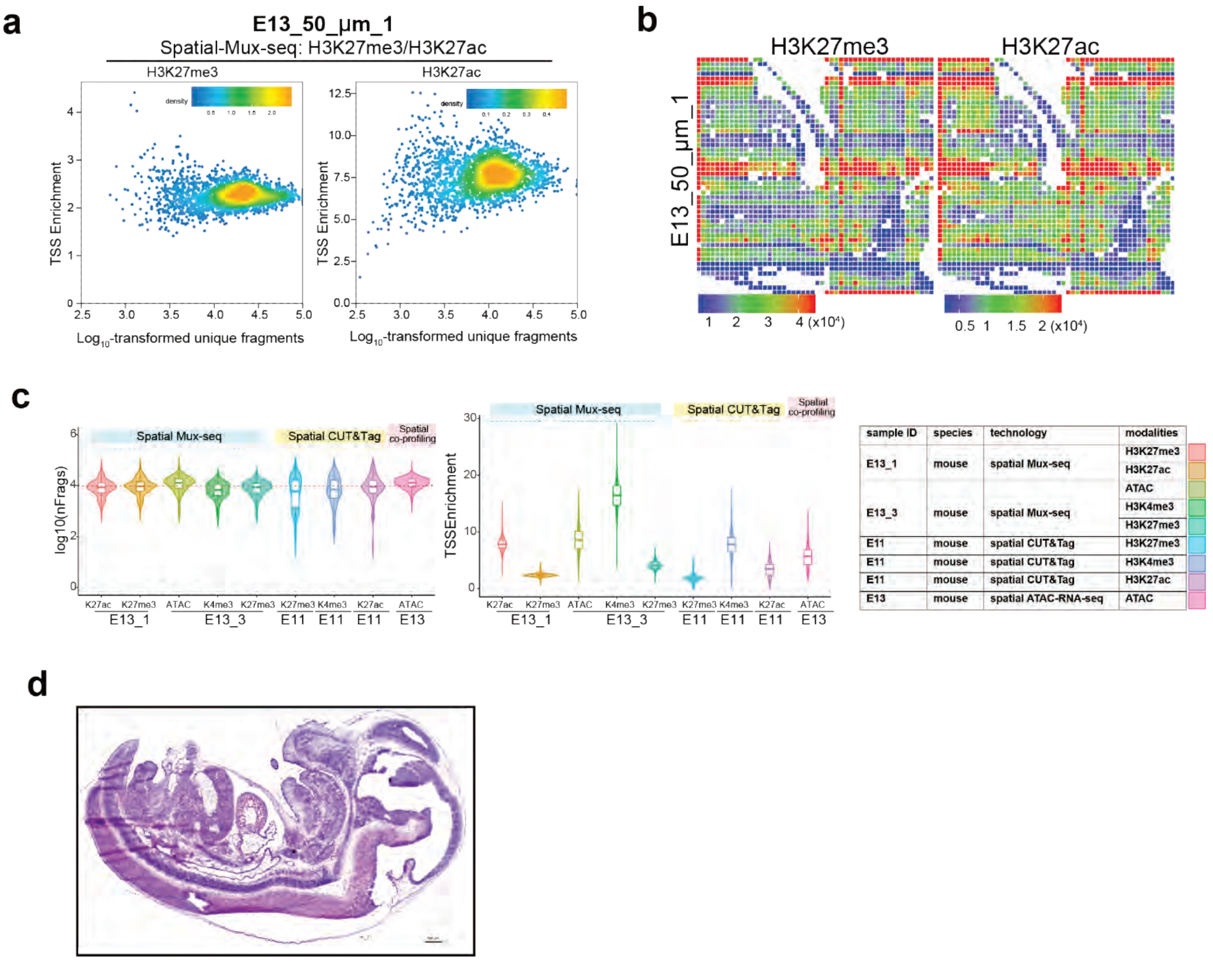


**Extended Data Fig. 2 | Quality control metrics for spatial-Mux-seq datasets.** sample: E13_50_μm_1. **a,** Scatterplots showing the TSS enrichment score vs unique nuclear fragments per pixel. **b,** Unique fragment counts in spatial-Mux-seq epigenome mapping of E13 mouse embryos (50 μm pixel size: H3K27me3 and H3K27ac). **c,** Comparison of number of unique fragments (left) and TSS enrichment (middle) between different spatial-Mux-seq (E13_50_μm_1: H3K27me3/H3K27ac or E13_50_μm_3: ATAC/H3K4me3/H3K27me3/RNA), spatial-CUT&Tag[11] (single modality: H3K4me3, H3K27me3, or H3K27ac), and spatial-ATAC-RNA-seq[4] (two modalities: ATAC/RNA) at a 50 μm resolution. Only epigenome modalities are compared. Fastq files were down-sampled to 50 million reads per sample for consistent comparison. **d,** H&E staining of E13 mouse embryo. Scale bar: 500 μm.

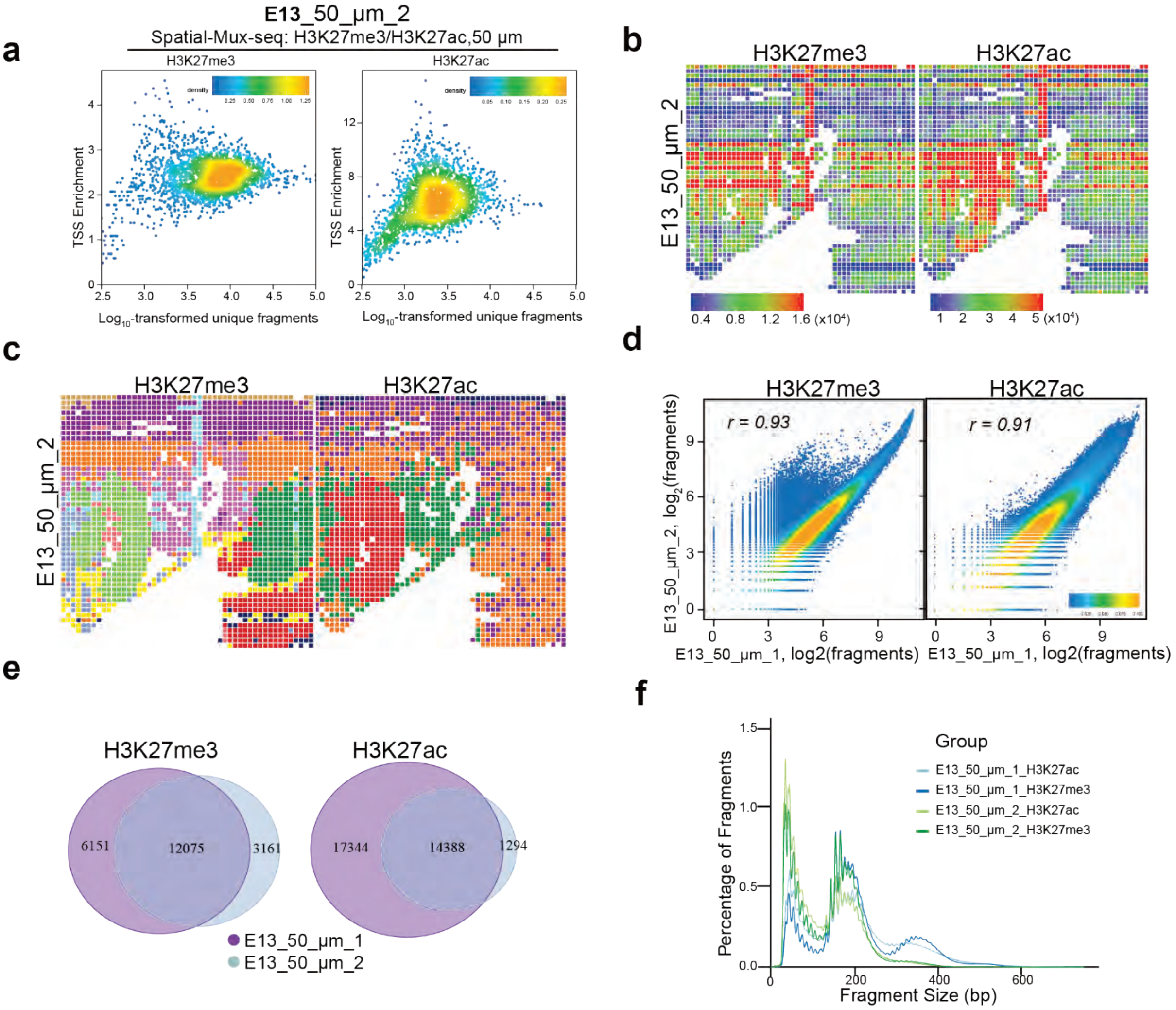


**Extended Data Fig. 3 | Reproducibility of Spatial-Mux-seq.** Spatial-Mux-seq profiling of a mouse embryo tissue section (sample: E13_50_µm_2). **a,** Scatterplots showing TSS enrichment score versus unique nuclear fragments per pixel. **b,** Unique fragment counts in spatial-Mux-seq epigenome mapping of sample E13_µm_2 (50 µm pixel size: H3K27me3 and H3K27ac). **c,** Unsupervised clustering analysis is performed, revealing the spatial distribution of clusters corresponding to H3K27me3 and H3K27ac mark. **d,** Reproducibility between two biological replicates (samples E13_µm_1 and E13_µm_2) is shown, comparing the data for H3K27me3 and H3K27ac histone modifications. **e,** Venn diagram shows the overlap of peaks from two different spatial-Mux-seq experiments (co-profiled H3K27me3/H3K27ac). **f,** Distribution of insert size for histone modification fragments in the spatial-Mux-seq datasets.

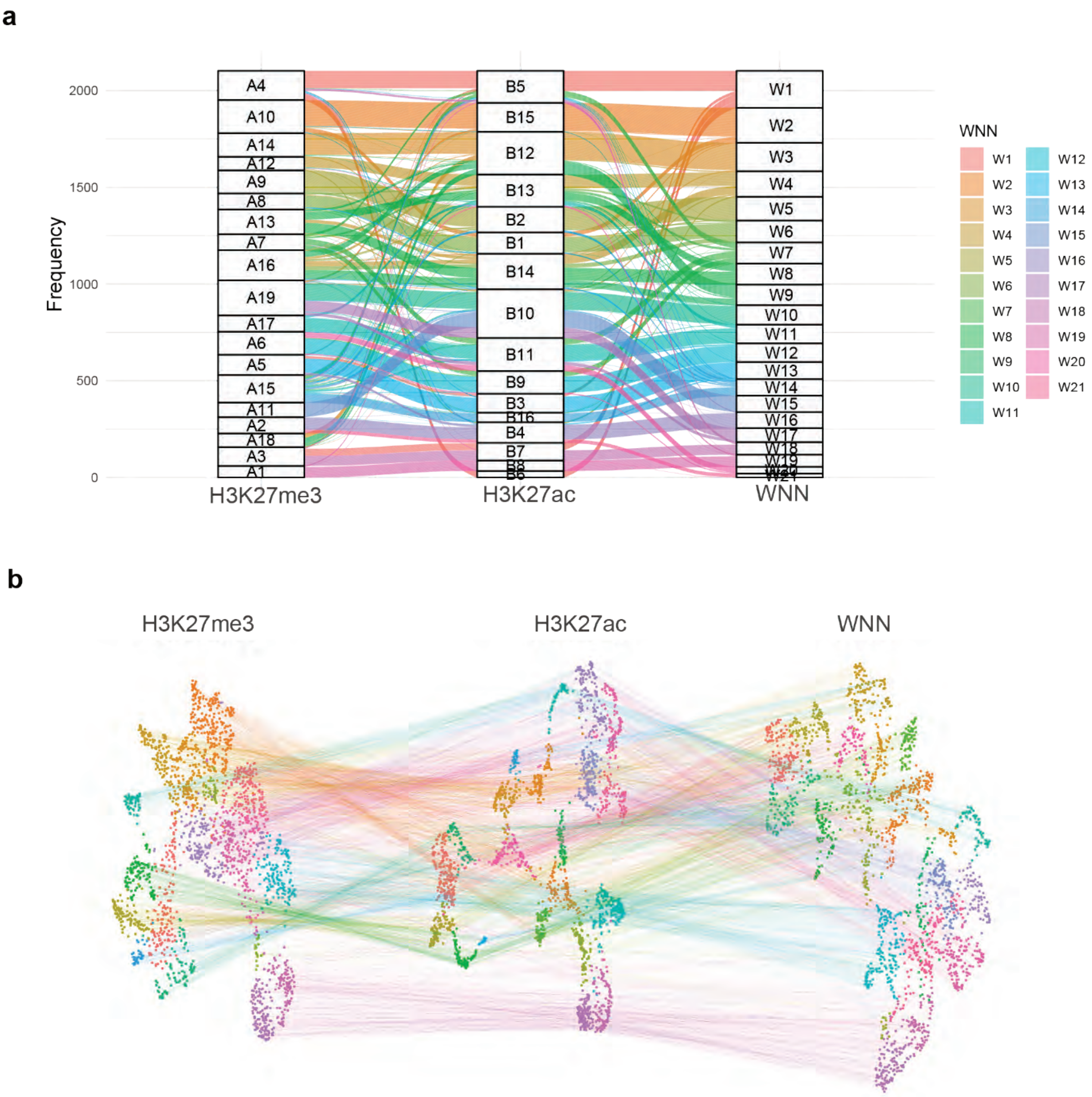


**Extended Data Fig. 4 | Alluvial diagram and UMAP embeddings of the spatial-Mux-seq data.** Sample: E13_50_μm_1. Alluvial diagram **(a)** and UMAP **(b)** of corresponding clusters from H3K27ac, H3K27me3, and WNN. The lines connect the same cells in the individual modalities.

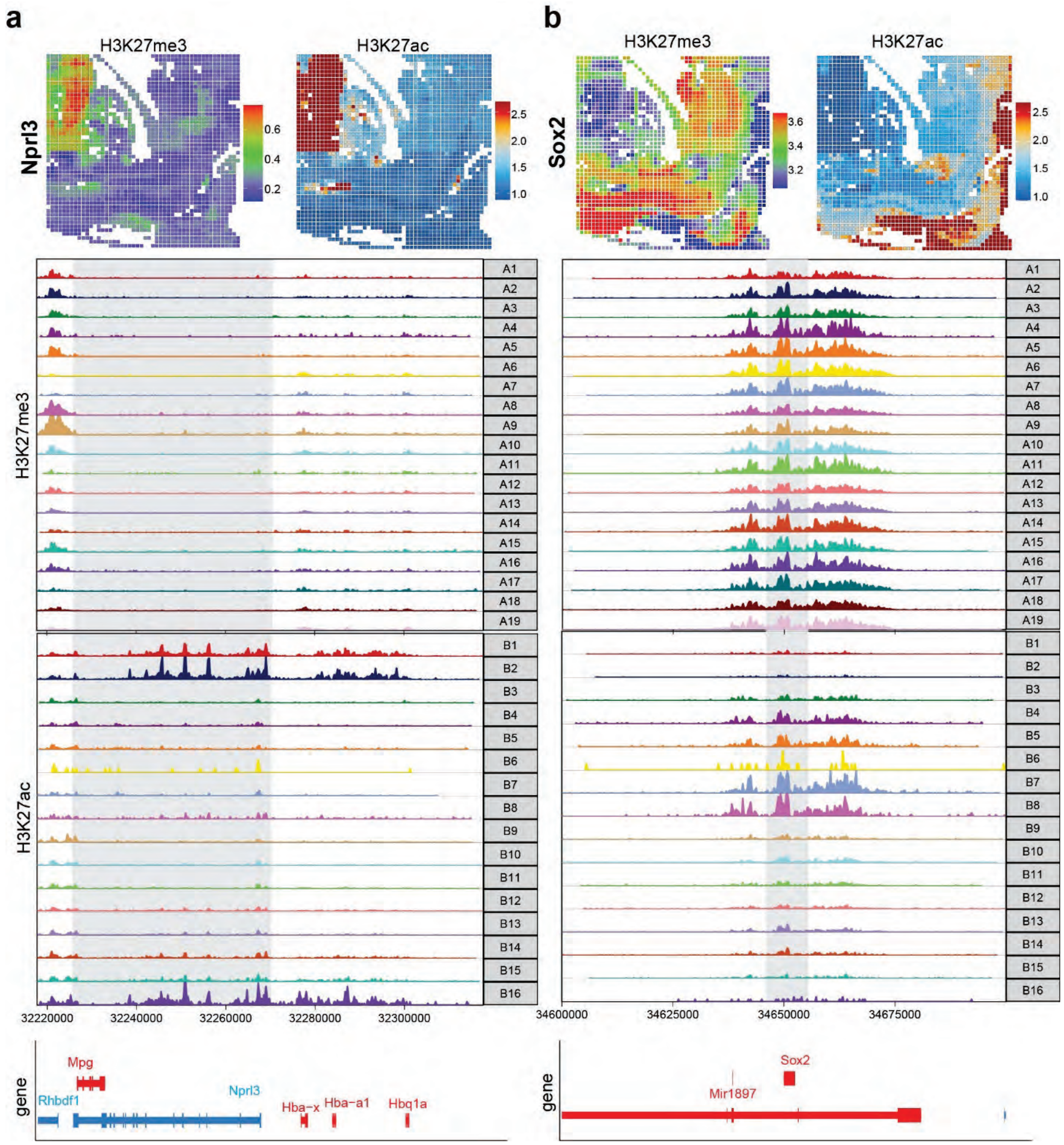


**Extended Data Fig. 5 | Spatial-Mux-seq (co-profiled H3K27me3/H3K27ac) mapping of marker genes in E13 mouse embryos.** The spatial mapping (top) and genome browser tracks (bottom) illustrate gene silencing marked by H3K27me3 and gene activity marked by H3K27ac modifications. Two marker genes are highlighted: *Nprl3* (**a**), representing gene silencing through H3K27me3, and *Sox2* (**b**), showcasing gene activity associated with H3K27ac modification.

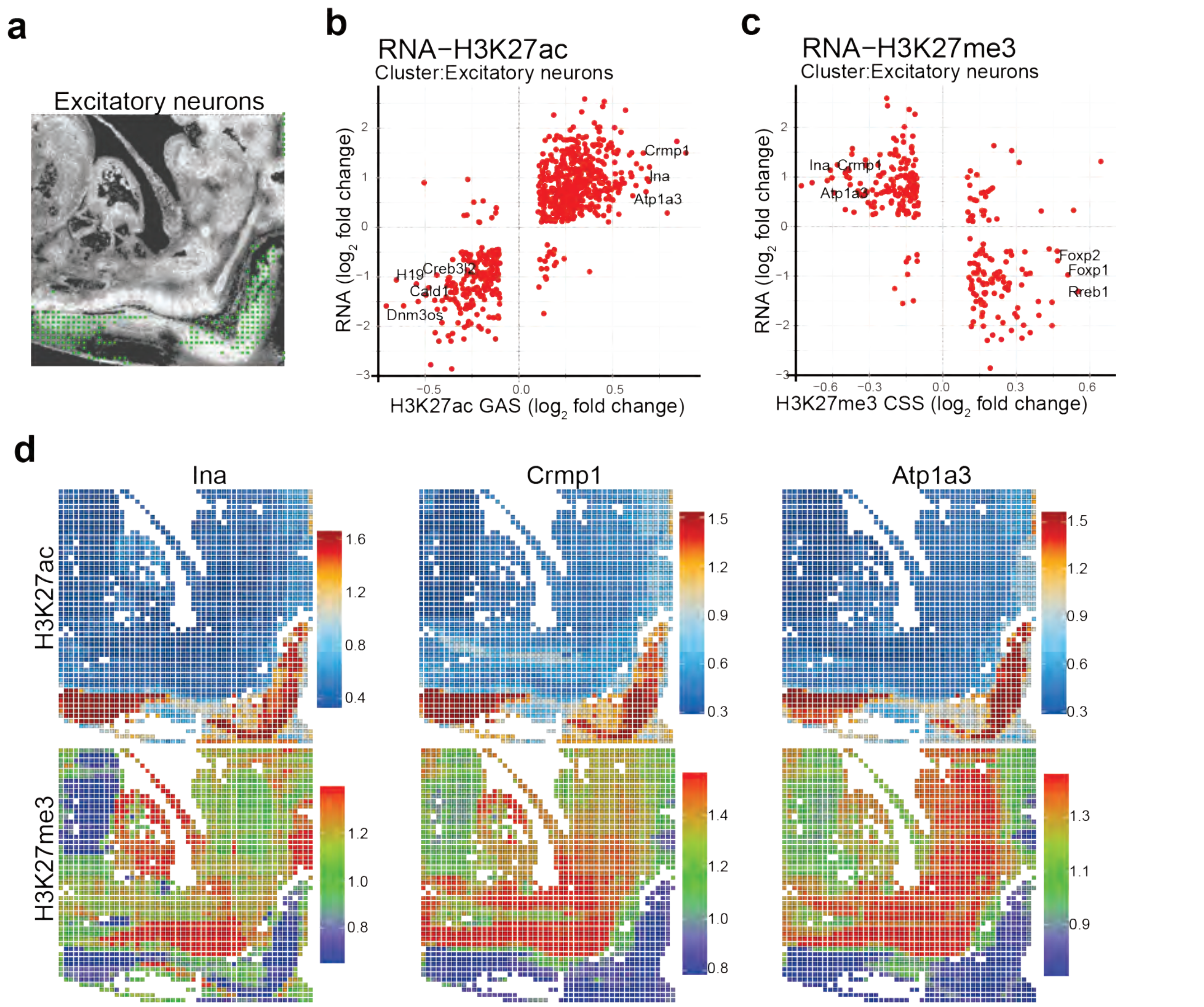


**Extended Data Fig. 6 | Spatial-Mux-seq (co-profiled H3K27me3/H3K27ac) mapping of marker genes in E13 mouse embryo. a,** Spatial mapping of excitatory neurons identified through label transferring, overlaid on a tissue section. Neuronal clusters are visualized with distinct patterns, emphasizing their spatial distribution within the embryo. **b,** Correlation of H3K27ac GAS and scRNA-seq data[14] in the cluster of excitatory neurons, highlighting the transcriptional activity associated with these regions. **c,** Correlation of H3K27me3 CSS and scRNA-seq data[14] in the cluster of excitatory neurons, emphasizing the gene silencing characteristics of these neurons. **d,** Heatmaps showing spatial mapping of marker genes associated with H3K27me3 and H3K27ac modifications, with variations in color intensity indicating differential expression and histone modification patterns across the embryo tissue.

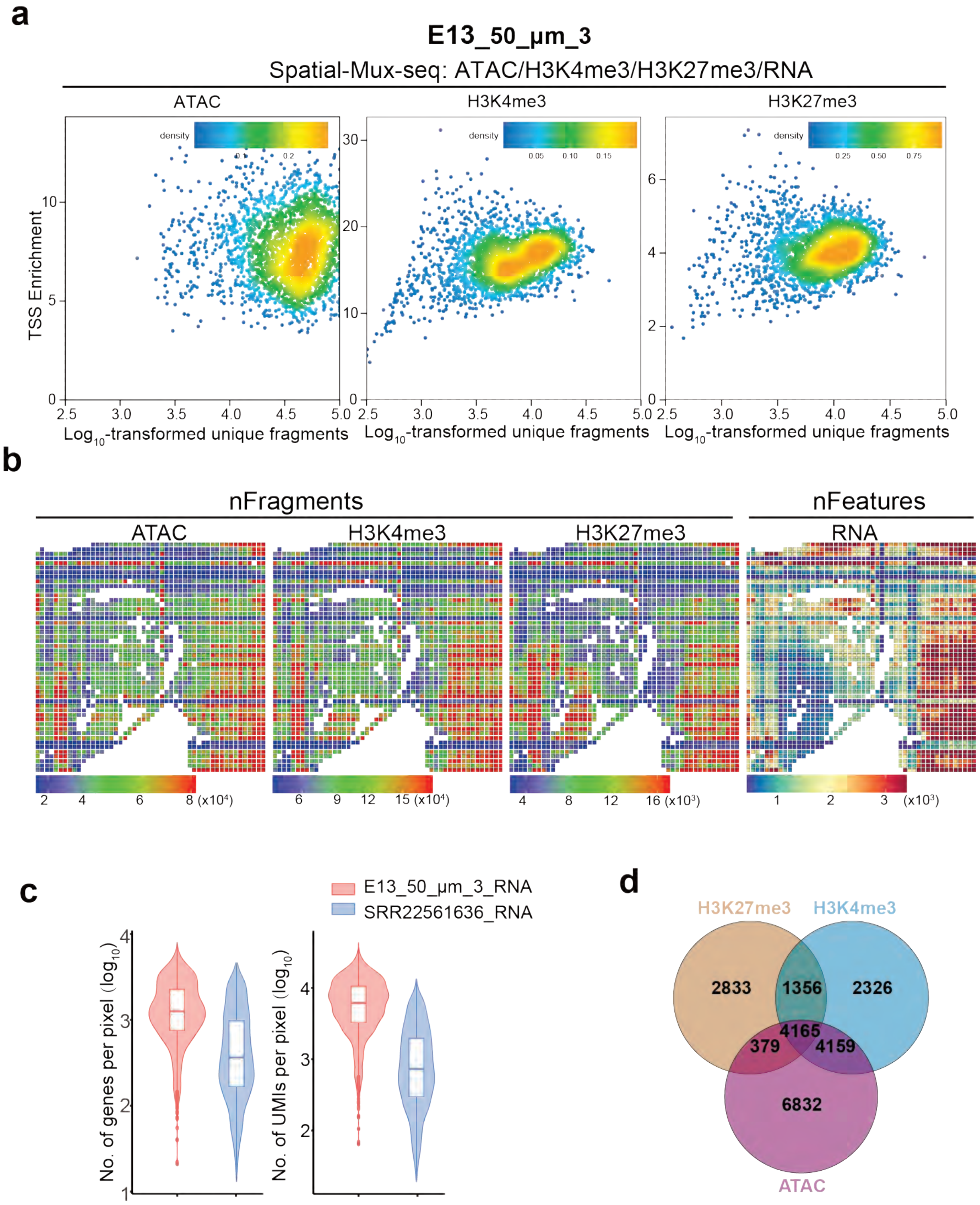


**Extended Data Fig. 7 | Quality control metrics for spatial-Mux-seq datasets.** Sample: E13_50_µm_3. **a,** Scatterplots showing the TSS enrichment score vs unique nuclear fragments per pixel for three modalities: ATAC, H3K4me3, and H3K27me3. **b,** Unique fragment or gene feature counts in spatial-Mux-seq four modalities mapping of E13 mouse embryos obtained with 50-µm pixel size. **c,** Gene and UMI Count Distribution: Comparison of RNA data between the E13_50_µm_3 sample and the RNA data from spatial-ATAC-RNA-seq[4] (SRR22561636). Fastq files were down-sampled to 50 million reads per sample for consistent comparison. **d,** Venn diagram showing the overlap of peaks identified from ATAC, H3K4me3, and H3K27me3 modalities. The liver clusters from all three modalities were extracted using the same pixel regions identified in the H3K27me3 data.

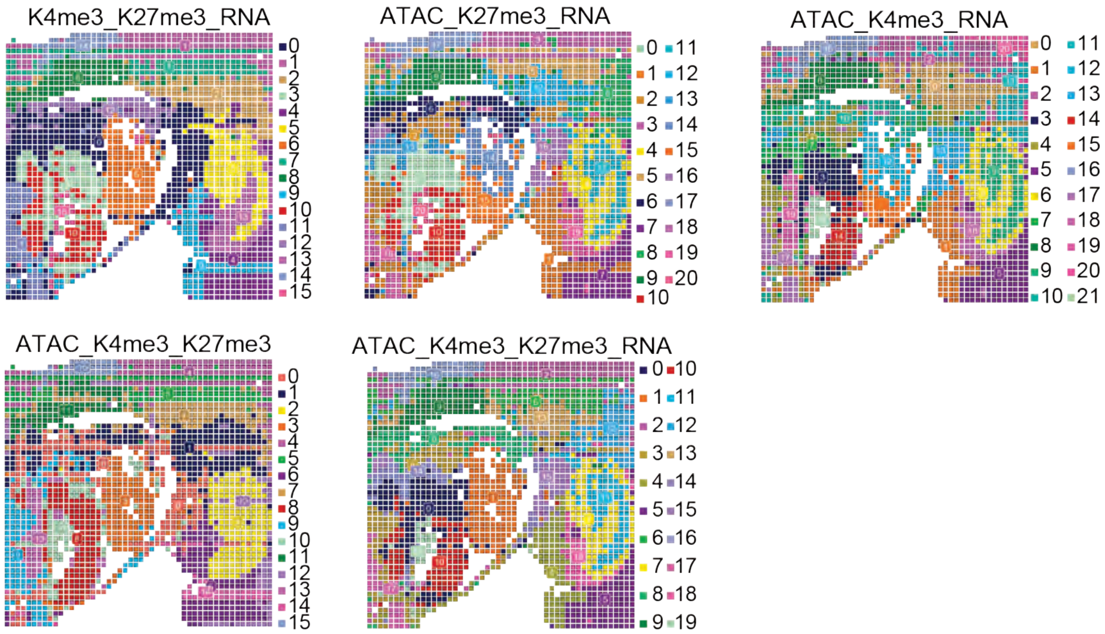


**Extended Data Fig. 8 | WNN Analysis of Co-Profiled Spatial-Mux-Seq Data (ATAC/H3K4me3/H3K27me3/RNA).** This figure presents the results of Weighted Nearest Neighbor (WNN) analysis on a spatial tissue section (sample: E13_50_µm_3). The analysis integrates different trimodal and quadrimodal data matrices, combining chromatin accessibility (ATAC), histone modifications (H3K4me3 and H3K27me3), and RNA expression profiles. The spatial distribution of the integrated clusters is visualized across the tissue map, with distinct regions colored to indicate specific cellular or tissue states identified by the WNN analysis.

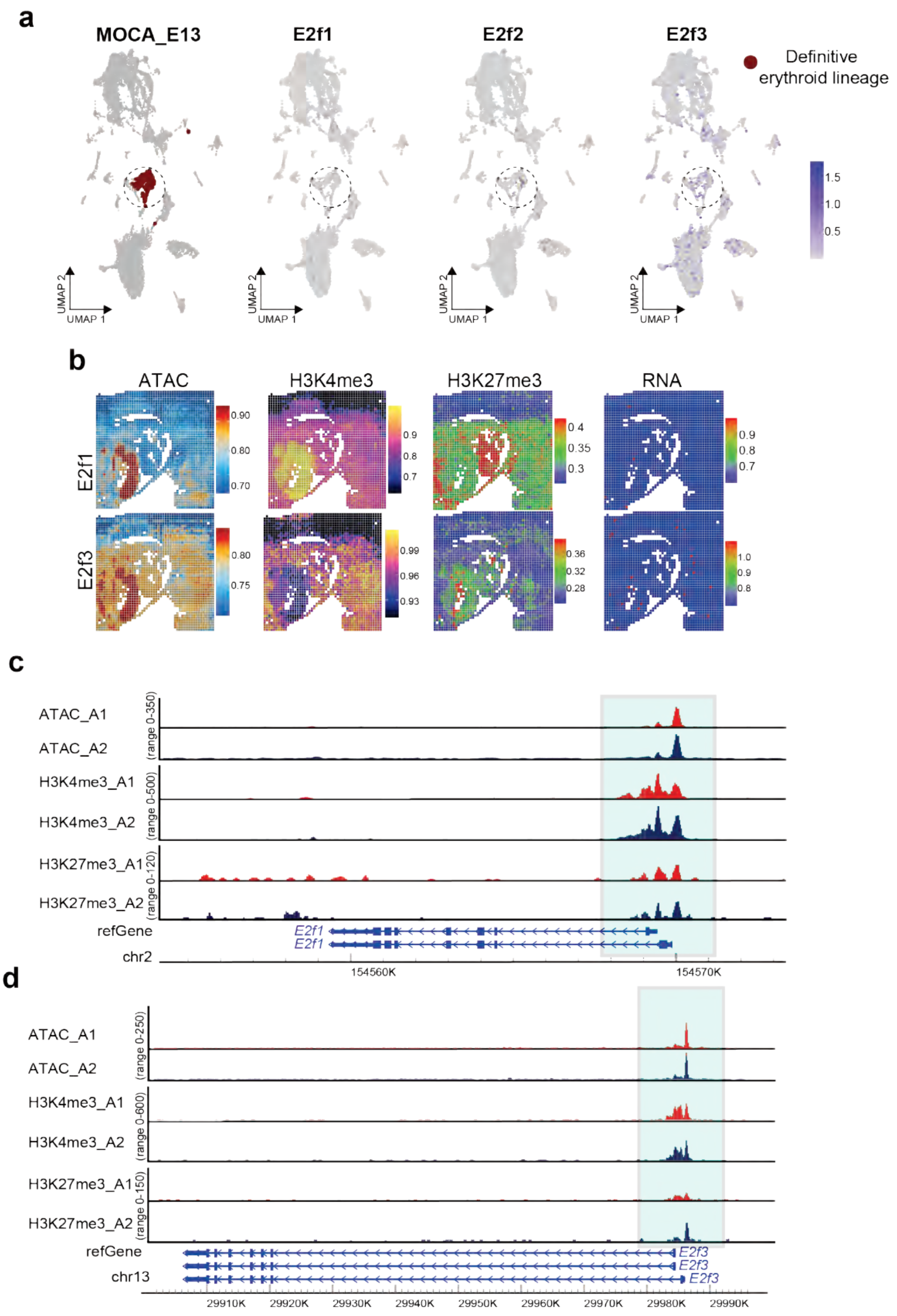


**Extended Data Fig. 9 | Spatial-Mux-seq (co-profiled H3K4me3/H3K27me3/ATAC/RNA) mapping of marker genes in E13 mouse embryo. a,** UMAP visualization of the scRNA-seq dataset[14] from mouse embryonic E13.5, highlighting the distribution of cells within the dataset. The expression patterns of the *E2f1-3* genes within the definitive erythroid lineage are specifically visualized, showcasing distinct clusters of gene activity. **b,** Spatial mapping of *E2f1* and *E2f3* gene expression across multiple modalities, including RNA, ATAC, H3K4me3, and H3K27me3. These heatmaps demonstrate the spatial distribution of gene activity and chromatin state modifications within the tissue section. **c-d,** Genome browser tracks of ATAC/H3K4me3/H3K27me3 marks for genes *E2f1* (**c**) and *E2f3* (**d**) in two liver clusters (A1 and A2) defined by spatial ATAC data.

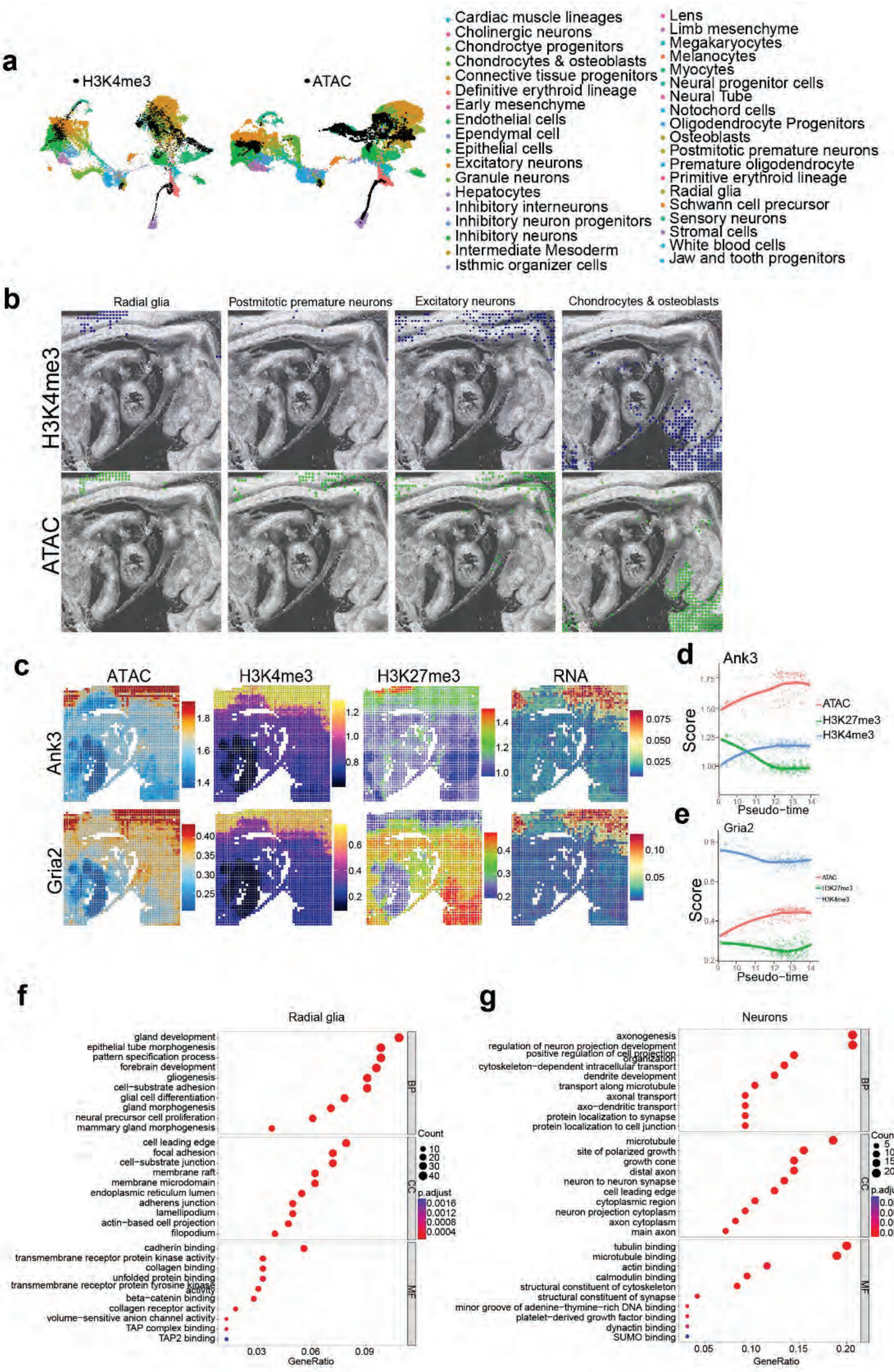


**Extended Data Fig. 10 | Spatial-Mux-seq (co-profiled H3K4me3/H3K27me3/ATAC/RNA) mapping of E13 mouse embryos a,** Spatial ATAC data and H3K4me3 data were integrated with scRNA-seq[14] from mouse embryo (E13.5). Unsupervised clustering of the combined data was colored by different cell types. **b,** Spatial mapping of selected cell types identified by label transferring from scRNA-seq to spatial H3K4me3 data or spatial ATAC data. **c,** Spatial mapping of *Ank3* and *Gria2* genes with RNA, ATAC, H3K4me3, and H3K27me3 modalities. **d-e,** Scatter plot showing scaled values of *Ank3* and *Gria2* ATAC, H3K4me3, and H3K27me3 score across pseudotime from radial glia to differentiated neurons. **f-g,** GO enrichment analysis for genes from radial glia (**f**) to differentiated neurons (**g**).

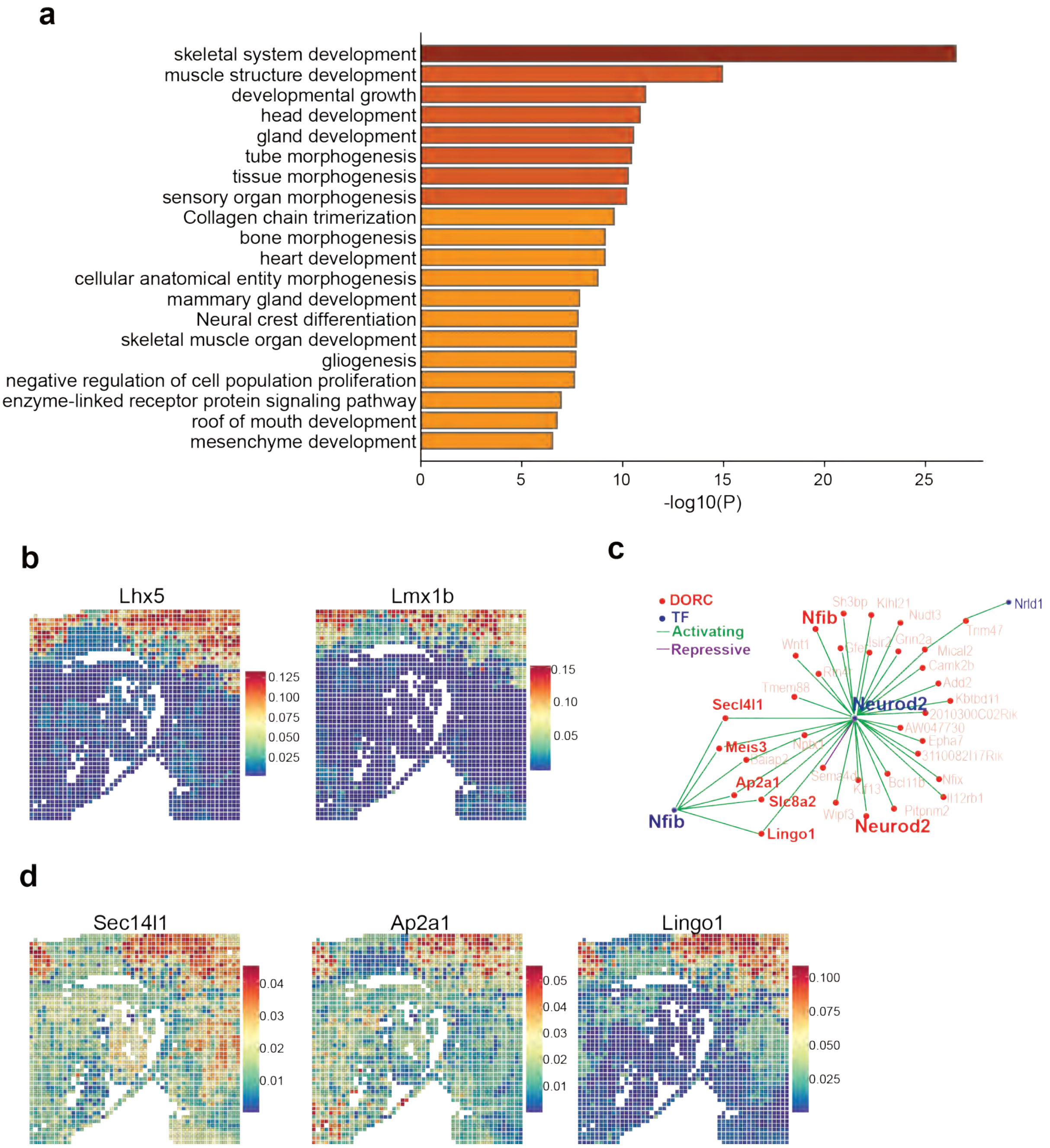


**Extended Data Fig. 11 | Integrative Spatial ATAC-RNA analysis identifies key regulatory modules associated with E13 mouse embryo development. a,** GO enrichment analysis for 411 DORC genes identified in **Fig. 2h**. **b,** Spatial mapping of *Lhx5* and *Lmx1b* genes with RNA modality. **c,** Gene regulatory network visualization of *Neurod2-Nfib* cascade. The width of the edges represents the regulation score. **d,** Spatial mapping of *Sec14l1, Ap2a1,* and *Lingo1* genes with RNA modality.

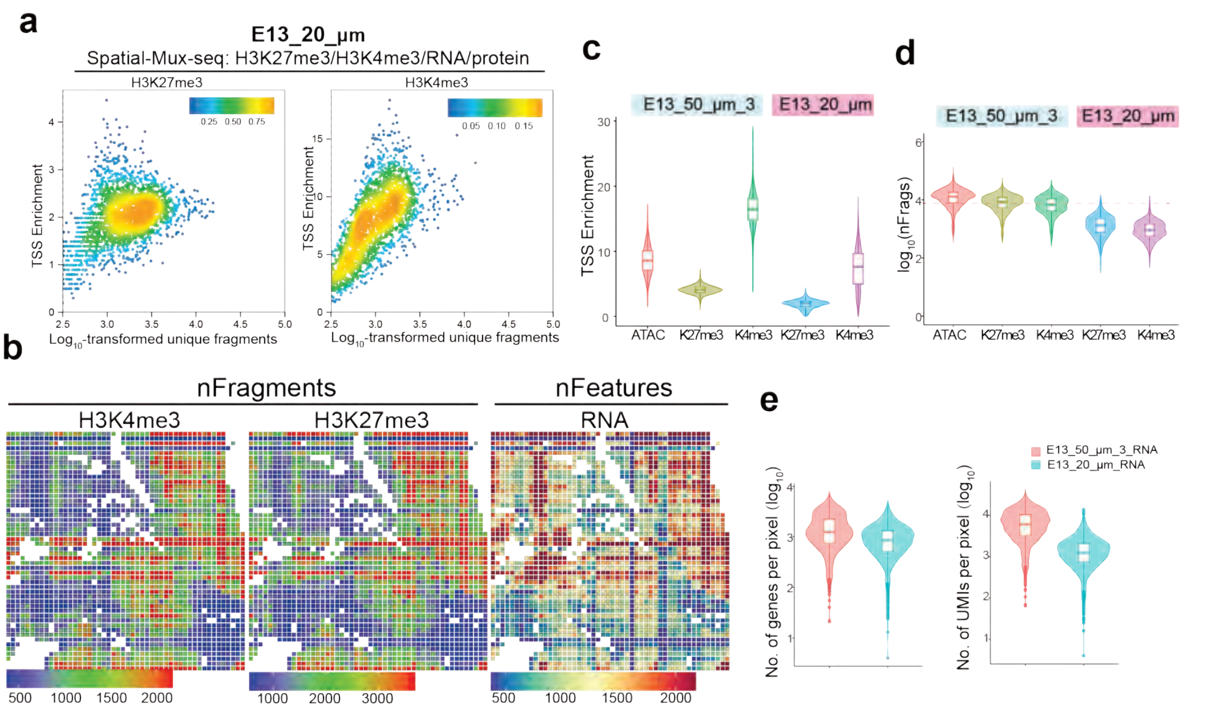


**Extended Data Fig. 12 | Quality control metrics for spatial-Mux-seq datasets.** Sample: E13_20_µm. **a,** Scatterplots showing the TSS enrichment score vs unique nuclear fragments per pixel for two modalities: H3K4me3 and H3K27me3. **b,** Unique fragment and gene feature counts in spatial-Mux-seq mapping of E13 mouse embryos obtained with 20-µm pixel size. **c-e,** The differences in data yield and quality between the different pixel resolutions. 50-µm and 20_µm pixel size devices were used to compare TSS enrichment (**c**), unique fragments (**d**), detected genes and UMI counts (**e**). Fastq files were down-sampled to match 50M read/sample. Samples analysed: E13_50_µm_3 and E13_20_µm.

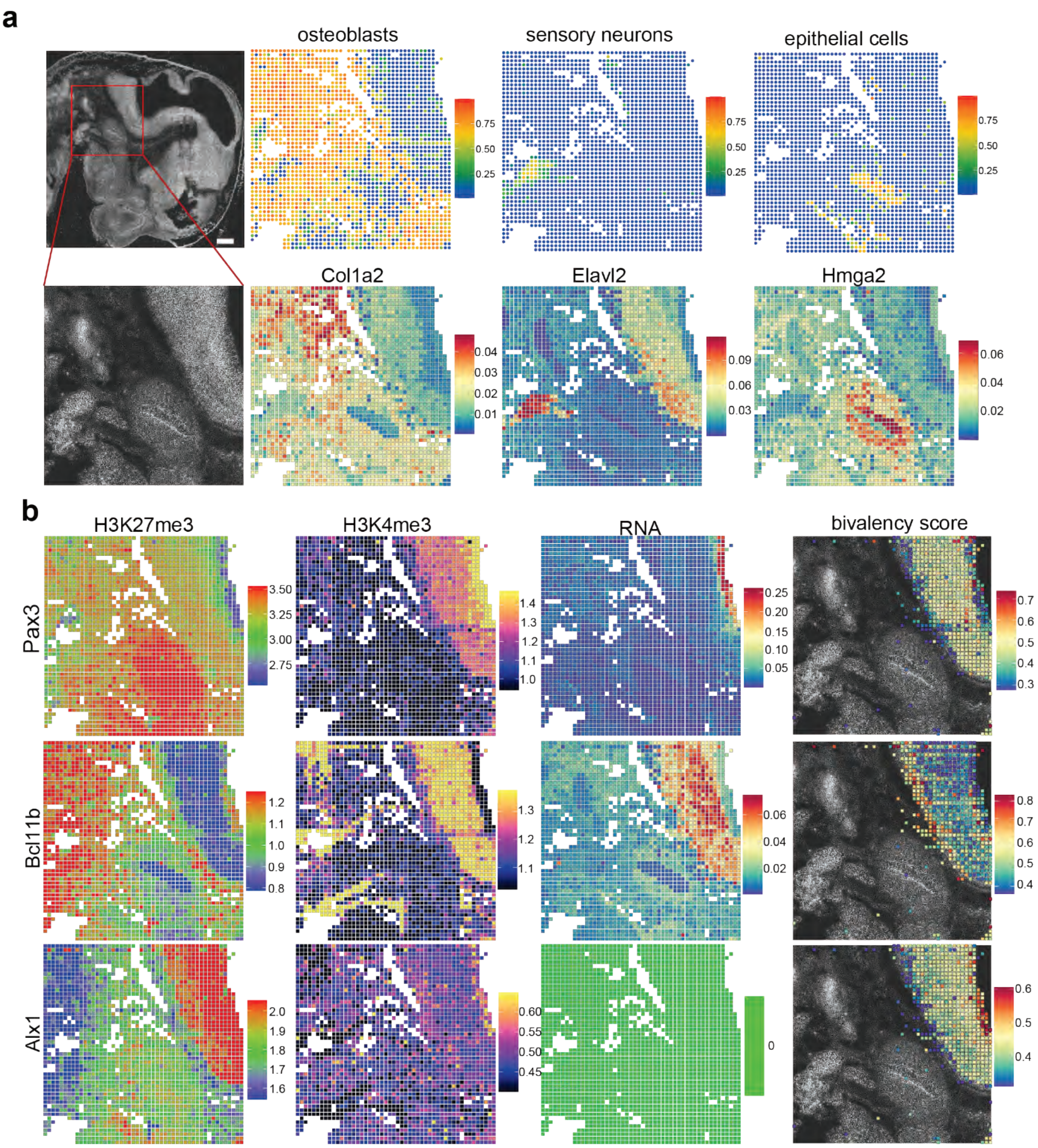


**Extended Data Fig. 13 | Spatial co-profiling of protein, RNA, H3K4me3, and H3K27me3 in mouse embryos. a,** Spatial RNA data were integrated with scRNA-seq[14] from E13.5 mouse embryos. This integration enabled the spatial mapping of specific cell types, including osteoblasts, sensory neurons, and epithelial cells within the embryonic tissue. The spatial patterns of marker genes of each cell type are performed with RNA modality. Red square indicates the region of interest (ROI). **b,** Spatial mapping of selected genes with RNA, H3K4me3, H3K27me3 and bivalency score. Scale bar: 500 μm.

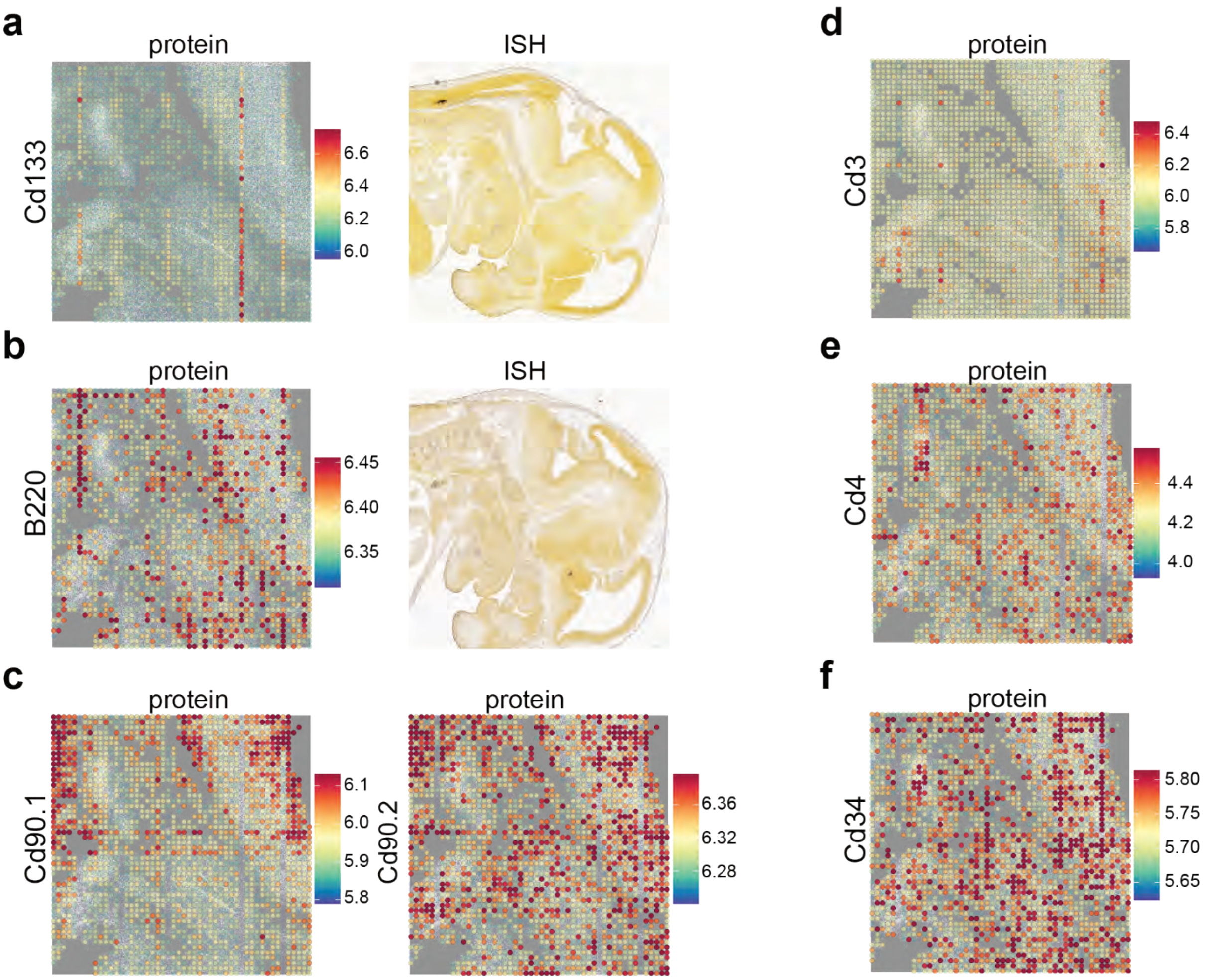


**Extended Data Fig. 14 | Spatial pattern of cell surface protein expression in mouse embryos.** The spatial distribution of cell surface markers, such as Cd133 (**a**) and B220 (**b**), was examined. The spatial pattern of these markers is depicted on the left. On the right, corresponding gene expression patterns are shown using *In Situ* Hybridization data sourced from the Allen Mouse Brain Atlas. **c-f,** Analysis of the distribution and localization of several cell surface proteins in mouse embryos, including Cd90 (**c**), Cd3 (**d**), Cd4 (**e**), and Cd34 (**f**). Cd90 (**c**): Represented by two distinct antibodies, Cd90.1 and Cd90.2, which recognize different epitopes or allelic variants of the Cd90 protein.

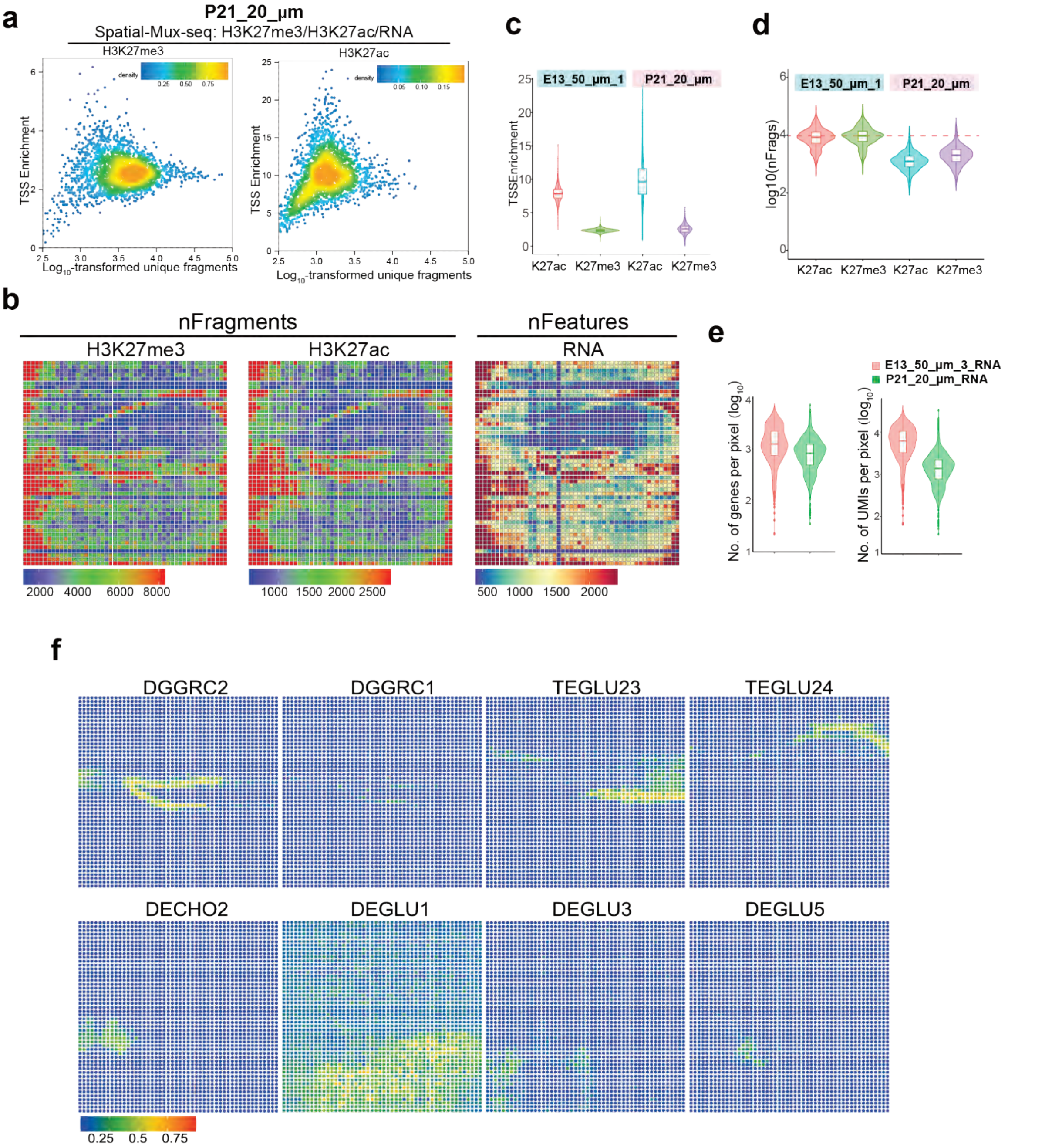


**Extended Data Fig. 15 | Quality control metrics for spatial-Mux-seq datasets.** Sample: P21_20_µm. **a,** Scatterplots showing the TSS enrichment score vs unique nuclear fragments per pixel for two modalities: H3K27me3 and H3K27ac. **b,** Unique fragment or gene feature counts in spatial-Mux-seq three modalities mapping of P21 mouse brain obtained with 20-µm pixel size. **c-d,** 50-µm and 20_µm pixel size devices were used to compare TSS enrichments (**c**) and unique fragments (**d**) of H3K27me3 and H3K4me3 modifications. The compared data are from E13_50_µm_1 and E13_20_µm samples. **e**, Comparison of gene counts and UMIs from 50-µm and 20_µm pixel size devices. The compared RNA data are from E13_50_µm_3 and E13_20_µm samples. Fastq files were down-sampled to match 50M read/sample. **f,** Integration of spatial RNA data with scRNA-seq data from mouse brain datasets enables high-resolution mapping of selected cell types, including dentate gyrus granule neuroblasts (DGGRC1), dentate gyrus granule neurons (DGGRC2), cornu ammonis excitatory neurons (TEGLU23 and TEGLU24), habenula cholinergic neurons (DECHO2), and thalamus excitatory neurons (DEGLU1, DEGLU3, and DEGLU5).

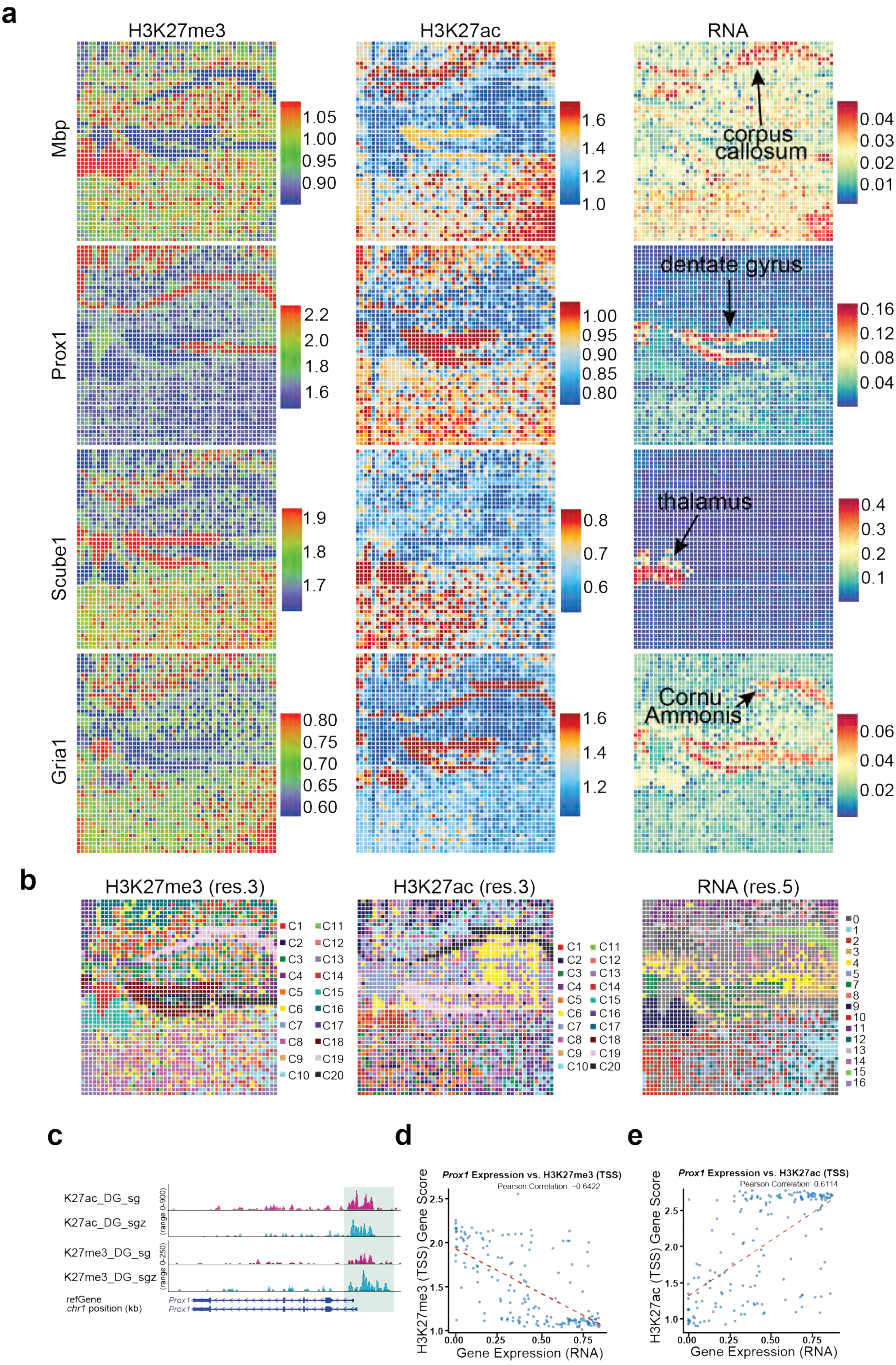
a
H3K27me3
H3K27ac
RNA
Mbp
Prox1
Scube1
Gria1
1.05
1.00
0.95
0.90
1.6
1.4
1.2
1.0
0.04
0.03
0.02
0.01
corpus callosum
2.2
2.0
1.8
1.6
1.00
0.95
0.90
0.85
0.80
0.16
0.12
0.08
0.04
dentate gyrus
1.9
1.8
1.7
0.8
0.7
0.6
0.4
0.3
0.2
0.1
thalamus
0.80
0.75
0.70
0.65
0.60
1.6
1.4
1.2
0.06
0.04
0.02
Cornu Ammonis
b
H3K27me3 (res.3)
H3K27ac (res.3)
RNA (res.5)
C1 C2 C3 C4 C5 C6 C7 C8 C9 C10 C11 C12 C13 C14 C15 C16 C17 C18 C19 C20
0 1 2 3 4 5 7 8 9 10 11 12 13 14 15 16
c
K27ac_DG_sg
K27ac_DG_sgz
K27me3_DG_sg
K27me3_DG_sgz
(range 0-900)
(range 0-250)
refGene
chr1 position (kb)
Prox1
Prox1
d
Prox1 Expression vs. H3K27me3 (TSS)
Pearson Correlation: -0.6422
H3K27me3 (TSS) Gene Score
Gene Expression (RNA)
e
Prox1 Expression vs. H3K27ac (TSS)
Pearson Correlation: 0.6114
H3K27ac (TSS) Gene Score
Gene Expression (RNA)

**Extended Data Fig. 16 | Spatial co-profiling of RNA, H3K27ac, and H3K27me3 in mouse juvenile brain. a,** Spatial mapping of selected genes with RNA, H3K27ac, and H3K27me3 modalities. **b,** Unsupervised clustering analysis and spatial distribution of each modality with different resolution from **Fig. 4a**: H3K27me3 (Resolution: 3), H3K27ac (Resolution: 3), and RNA (Resolution: 5). **c,** Genome browser tracks of *Prox1* gene in clusters DG-sg and DG-sgz. **d-e,** Pearson correlation between *Prox1* expression and histone mark H3K27me3 (**d**) or H3K27ac (**e**) gene scores. The gene scores are derived based on the gene model surrounding the transcription start site (TSS).

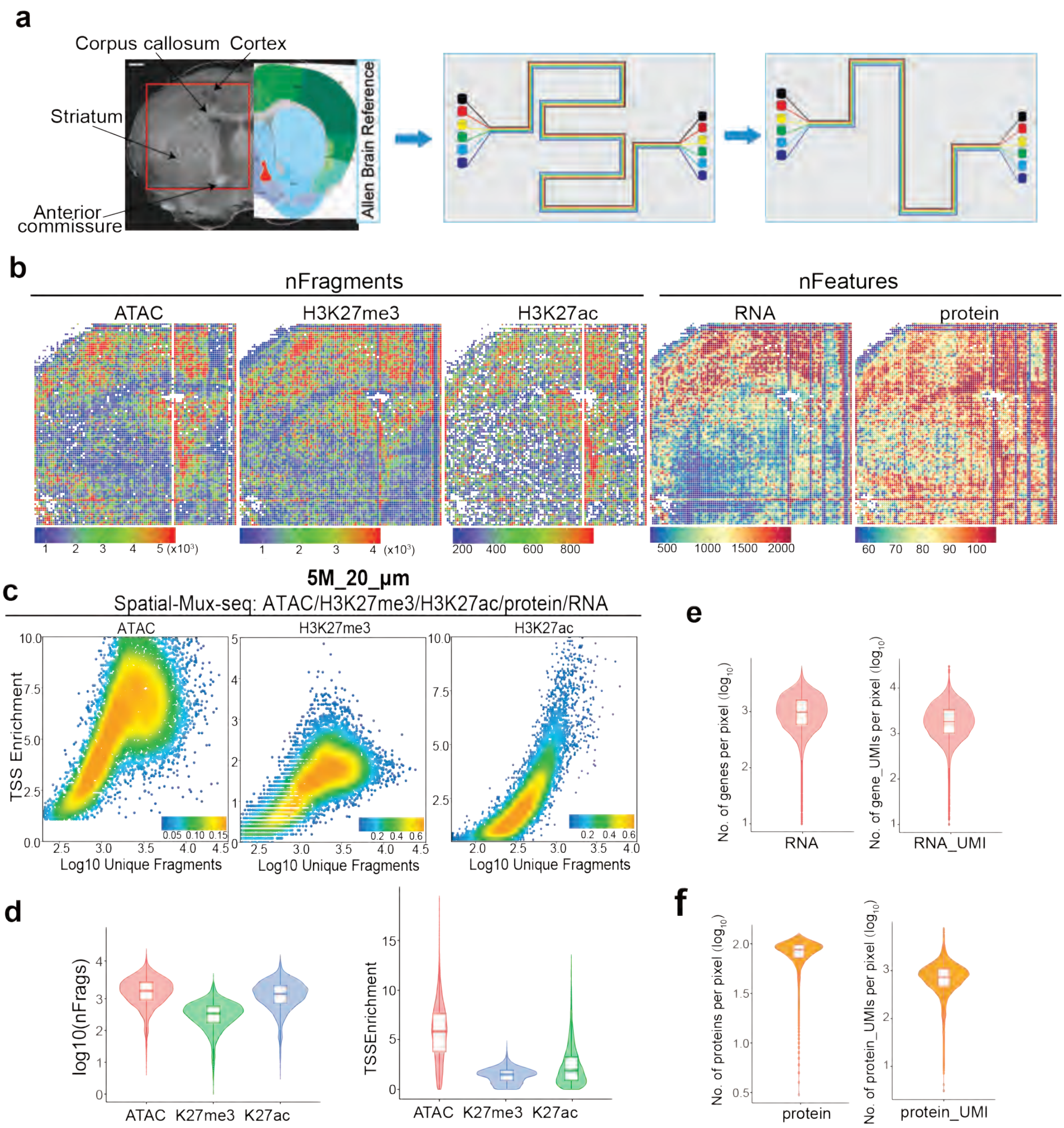


**Extended Data Fig. 17 | Quality control metrics for spatial-Mux-seq datasets.** Sample: 5M_20_µm. **a,** Spatial-Mux-seq profiling of ATAC, H3K27me3, H3K27ac, RNA, and proteins from EAE mouse brain section. Left: tissue scanning of the region of interest, aligned with the region annotation of a corresponding section from Allen Mouse Brain Atlas (P56). Middle and right: 20-µm-microfluidic device with 100x100 pixels. Two-time spatial barcodes ($A_{1\text{-}100}$ and $B_{1\text{-}100}$) were sequentially flowed over tissue section. **b,** Unique fragments, gene feature counts, and protein counts in spatial-Mux-seq mapping of five months mouse brain obtained with 20-μm pixel size. **c,** Scatterplots showing the TSS enrichment score vs unique nuclear fragments per pixel for three modalities: ATAC, H3K27me3 and H3K27ac. **d,** Violin plots of unique fragments and TSS enrichment values of ATAC, H3K27ac, and H3K27me3. **e,** Violin plots of gene counts and gene UMIs distribution. **f,** Violin plots of protein counts and protein UMIs distribution. Scale bar: 500 μm.

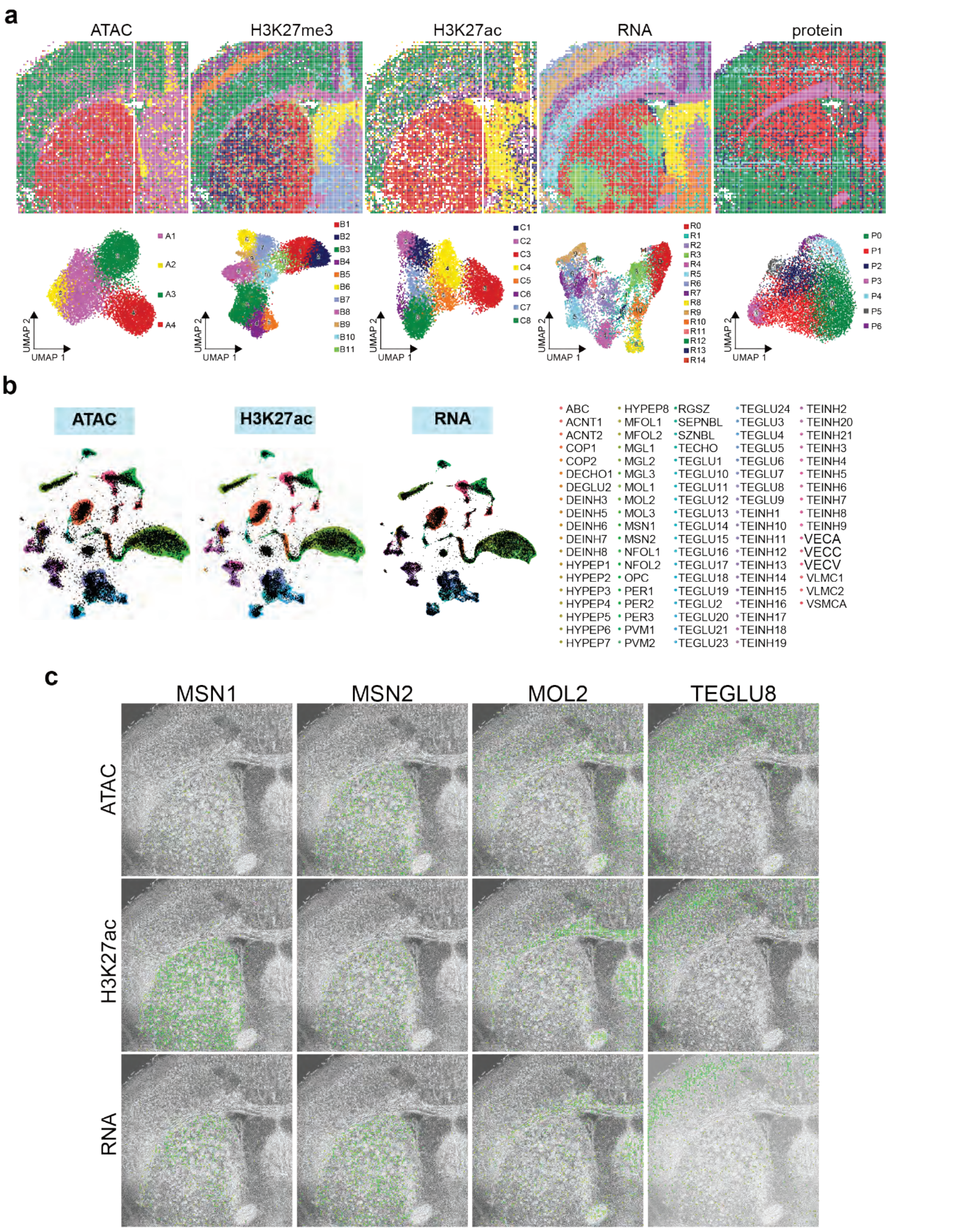


**Extended Data Fig. 18 | Spatial co-profiling of proteins, mRNA, H3K27me3, and H3K27ac in a EAE mouse brain.** Sample: 5M_20_μm. **a,** Spatial distribution and UMAP embeddings of unsupervised clustering analysis of ATAC (An), H3K27me3 (Bn), H3K27ac (Cn), RNA (Rn), and proteins (Pn) with five months old EAE mouse brain sample (20 μm pixel size). **b,** Spatial ATAC, H3K27ac, and RNA data were integrated with scRNA-seq[38] from mouse brain. Spatial mapping of cell types identified by label transfer from scRNA-seq to ATAC (top), H3K27ac (middle), and RNA (bottom). MSN: medium spiny neurons. MOL: mature oligodendrocytes 2. TEGLU: Telencephalic Glutamatergic Neurons.

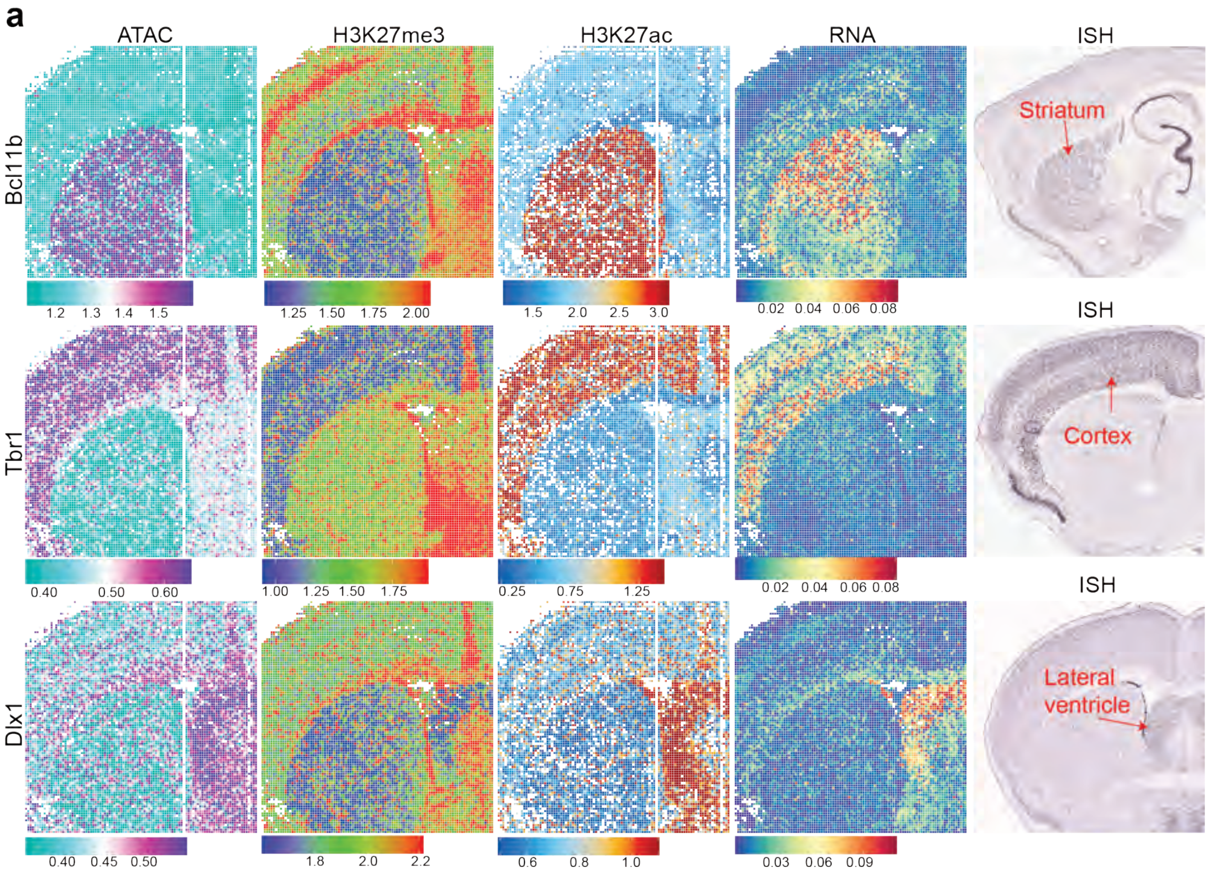


**Extended Data Fig. 19 | Spatial co-profiling of ATAC, H3K27me3, H3K27ac, and RNA of selected genes for 5M-old mouse brain.** Spatial mapping marker genes: *Bcl11b*, *Tbr1*, and *Dlx1* by all four modalities. *In Situ* hybridization (right) of corresponding genes were from the Allen Mouse Brain Atlas. Arrows indicate the high expression region of marker genes.

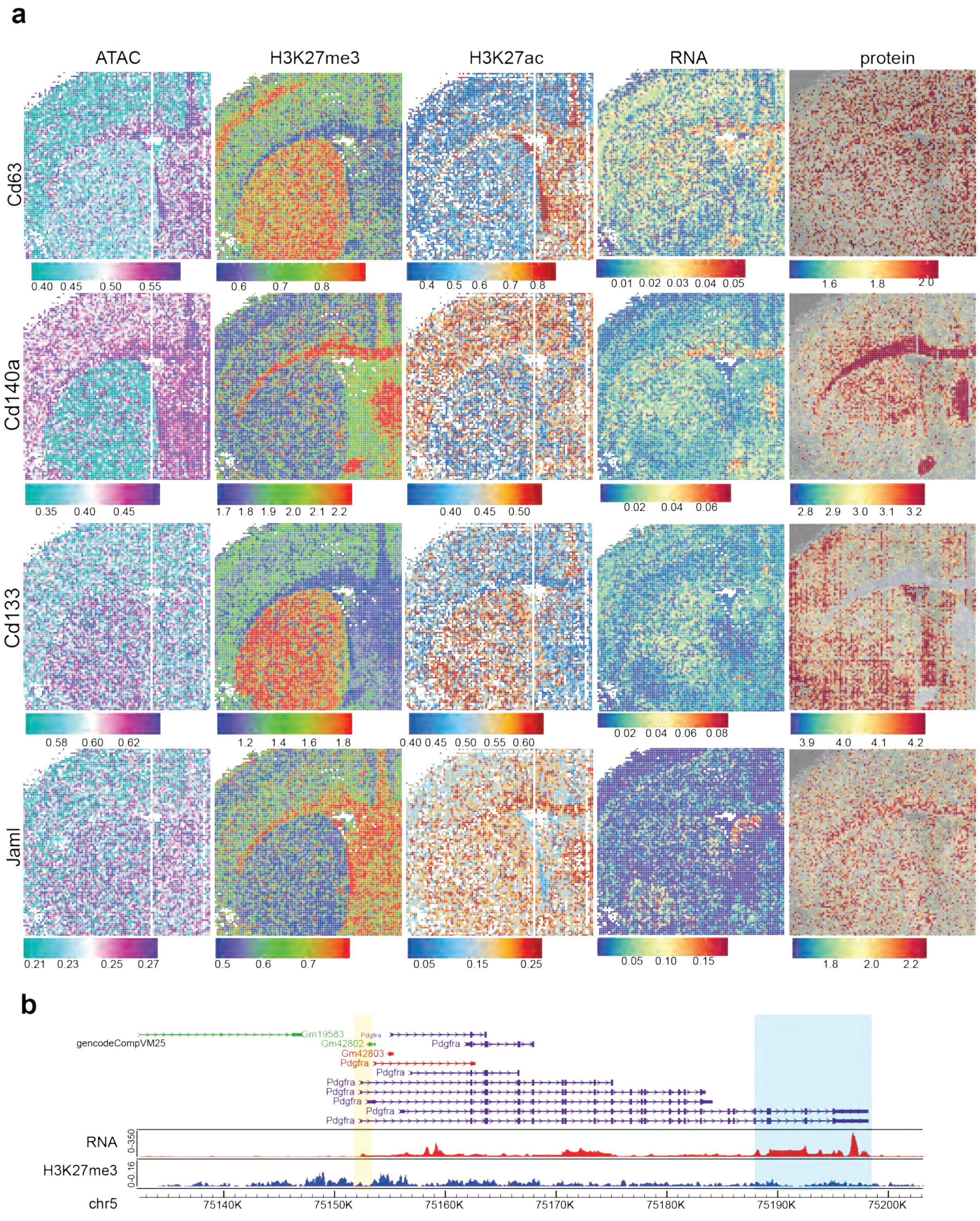


**Extended Data Fig. 20 | Spatial co-profiling of ATAC, H3K27me3, H3K27ac, protein, and RNA of selected genes for 5M-old mouse brain. a,** Spatial mapping marker genes: *Cd63*, *Cd140a*, *Cd133*, and *Jaml* by all five modalities: ATAC, H3K27me3, H3K27ac, protein, and RNA. **b,** Genome browser tracks of *Cd140a (Pdgfra)* gene expression and H3K27me3 signal in corpus callosum defined by spatial H3K27me3 data (cluster B8 from Extended Data Fig.18a).
